# Dynamic genetic adaptation of *Bacteroides thetaiotaomicron* murine gut colonization

**DOI:** 10.1101/2022.02.23.481734

**Authors:** Manjing Zhang, Megan Kennedy, Orlando DeLeon, Jacie Bissell, Florian Trigodet, Karen Lolans, Sara Temelkova, Katherine T. Carroll, Aretha Fiebig, Adam Deutschbauer, Ashley M. Sidebottom, Chris Henry, Phoebe A. Rice, Joy Bergelson, Eugene B. Chang

## Abstract

To understand how a bacterium ultimately succeeds or fails in adapting to a new environment, it is essential to assess the temporal dynamics of its fitness over the course of colonization. The mammalian gut, into which exogenous microorganisms are regularly introduced, represents a biologically and clinically relevant system to explore microbial adaptational processes. In this study, we introduce a human-derived commensal organism, *Bacteroides thetaiotaomicron*, into the guts of germ-free mice to 1) determine whether the genetic requirements for colonization shift over time and, if so, 2) characterize the biological functions required for microbial survival at different points of colonization. The results of a high-throughput functional genetics assay (BarSeq), transcriptomics, and metabolomics converge on several conclusions. First, adaptation to the host gut occurs in distinct stages. We observed drastic changes in gene usage during the first week, shifting from high expression of amino acid biosynthesis to polysaccharide utilization genes. These changes were sustained thereafter, except for the continued upregulation of a single polysaccharide utilization locus responsible for the degradation of raffinose-family oligosaccharides rich in the standard chow diet fed to our mice. Spontaneous mutations in wildtype *Bt* also evolve around this locus, highlighting the importance of efficient carbohydrate metabolism in long-term persistence within a monoassociated gut. To improve microbiome-based therapies, it will be important to appreciate and meet the distinct needs of the organism during each stage of colonization.

**Importance:** Microbes regularly disperse across and adapt to new environments and ecological niches. A clinically significant microbial niche home to trillions of microbes is the mammalian gut. Temporal processes of microbial adaptation over the course of gut colonization are poorly understood on a genetic, transcriptional, and metabolite level. In this study, we leverage a three-pronged approach to characterize gut colonization as a dynamic process with shifting genetic determinants of microbial fitness. This study sheds light on host colonization by *Bacteroides thetaiotaomicron,* an organism that is prevalent and dominant across healthy human microbiomes, and not only identifies key pathways involved in colonization, but determines the timing of *when* these pathways are most vital to colonization success. By demonstrating that the key determinants of colonization success in the gut change over time, the results of this study highlight the importance of considering ecological dynamics in developing more effective microbiome-based therapies.

## Introduction

Fast adaptation is paramount to the survival of any species undergoing an environmental transition. For microbial taxa, which are frequently and rapidly dispersed across dramatically different habitats and microenvironments, processes of local adaptation may arise as primary determinants of microbial colonization success and resulting biogeography (1). The mammalian gut is an environment regularly bombarded with a diverse array of exogenous microorganisms. As such, it represents a biologically and clinically relevant system to explore rapid microbial adaptational processes.

Canonically, adaptation occurs by way of genetic evolutionary mechanisms: genetic variants have differential fitness in the new environment, which establishes variation in survival and reproduction, and ultimately leads to shifts in gene frequencies over time. Over shorter timescales, organisms may adapt by altering plastic gene expression profiles to optimize functional characteristics including resource use, growth strategies, and resistance to environmental stressors (2–4). Both processes fundamentally reflect changes in fitness over time. To understand how a microbe ultimately succeeds or fails in adapting to a new environment, it is therefore essential to assess the temporal dynamics of its fitness over the course of colonization.

In the context of host-microbe interactions, adaptational processes play out within a dynamic host environment, in which feedbacks between microbe and host can have significant consequences for the fitness and survival of both (5). Understanding genetic and transcriptional changes of exogenous microbes upon entering a novel host system is a critical step toward elucidating the shifting landscapes created because of this feedback circuit. While previous work has evaluated the genetic requirements for colonization success across a variety of microbes in the guts of both conventionally raised and germ-free mice, these studies have generally assessed fitness at only a single timepoint, and therefore yield an incomplete picture of the temporal processes of adaptation and colonization (6–8). Moreover, although host transcriptional responses to microbial colonization are regularly documented, few studies have comprehensively assessed microbial gene expression profiles over the course of colonization (9–11).

In the following experiments, we introduce a human-derived commensal organism, *Bacteroides thetaiotaomicron* (*Bt*), into the guts of germ-free mice to 1) determine whether the genetic requirements for colonization shift over time and, if so, to 2) characterize the biological functions required for microbial survival at different timepoints of colonization. Use of a germ-free monocolonization model allows us to reduce the staggering complexity of the gut microbial ecosystem into experimentally tractable and readily interpretable components: here, we definitively outline population-level microbial colonization dynamics and the host-microbe interactions that drive them. Using this germ-free model as a baseline, future work can further establish the distinct contributions of microbe-microbe interactions and other emergent community-level properties to community assembly and colonization dynamics. To identify the microbial genes important for fitness in the context of gut monocolonization, we combine two complementary unbiased approaches: transcriptomics (RNA-seq), which reveals global gene usage patterns, and a functional genetics approach (BarSeq (8, 12)) to assess fitness consequences of gene disruptions at a global scale over the course of colonization. Finally, we evaluate spontaneous evolution of wildtype (WT) *Bt* in the gut to survey natural population-level fitness dynamics. Our results indicate that adaptation to the host gut occurs in distinct stages. During the earliest stage of colonization, genes involved in amino acid and vitamin biosynthesis are upregulated and, in some cases, play essential roles in survival of *Bt*. By the end of the first week, expression of these genes is downregulated, and expression of carbohydrate metabolism genes peaks. These levels are sustained for the rest of the two-week experimental period, except for the continued upregulation of a single polysaccharide utilization locus (PUL) responsible for the degradation of raffinose-family oligosaccharides (RFOs) rich in the standard chow diet fed to our mice. Spontaneous mutations in WT *Bt* also evolve around this locus, highlighting the importance of efficient carbohydrate metabolism in long-term persistence within a monoassociated gut.

These experiments lay the groundwork for future delineation of shifting colonization pressures in various host backgrounds and microbiome compositions. We expect that these insights into the temporally dynamic stresses that microbes must overcome to colonize and persist in the gut will prove invaluable to our understanding of microbial adaptation and the development of microbiome-based therapies.

## Results

### Both transcriptional and genetic fitness determinants shift over the course of Bt colonization and persistence

To evaluate global transcription during colonization, we introduced WT *Bt* into germ-free (GF) C57Bl/6 mice and collected cecal contents at Days 1, 7, and 14 after colonization (Fig. 1A). After rRNA and host RNA depletion, the bacterial RNA samples were sequenced and compared in a pairwise fashion across D1-D7 and D7-D14. In parallel, to assess functional genetic requirements during colonization, we introduced a rich library of randomly barcoded Tn insertion (RB-Tn) mutants of *Bt* into four different cohorts of germ-free C57Bl/6 mice and collected daily fecal samples (Fig. 1A). Amplification and sequencing of the transposon barcodes reveals the relative abundance of each mutant in the library at each timepoint.

**Figure 1:**
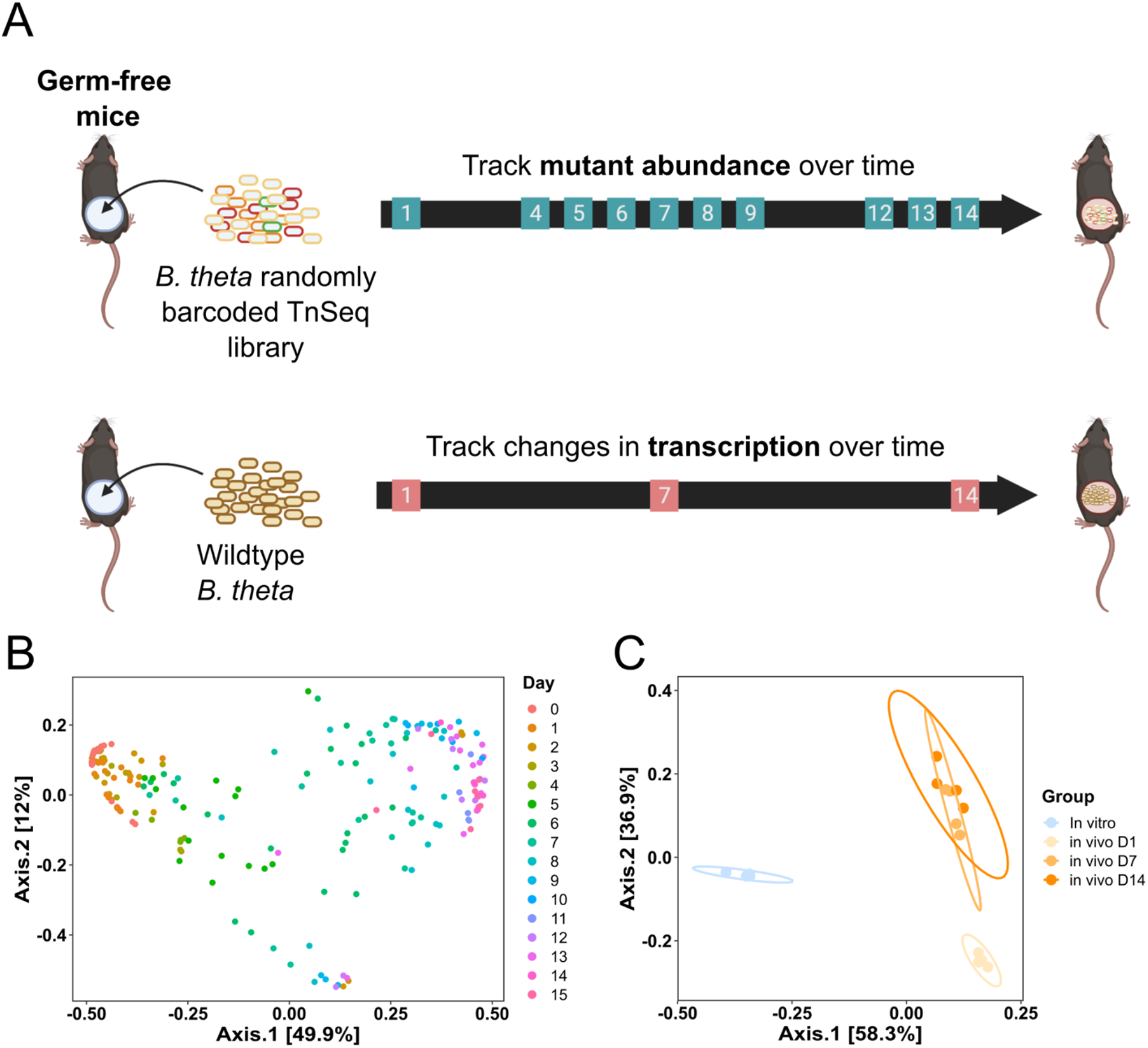
Bt gene expression and genetic fitness determinants shift over the course of colonization and persistence. A) Experimental scheme of functional genetics (top) and transcriptomics (bottom) experiments in germ-free mice. B) Principal Coordinates Analysis (PCoA) using Bray-Curtis dissimilarity on the relative abundance of RB-Tn mutant strains within each mouse, colored by experimental day. Four different cohorts of mice (n=3-5 per cohort) are represented. Samples clustered significantly by day (p = 0.0001, R^2^ = 0.457, PERMANOVA), experimental run (p = 0.0001, R^2^ = 0.092), and mouse ID (p = 0.0001, R^2^ = 0.135). C) PCoA using Bray-Curtis dissimilarity on the expression profiles of Bt within cecal samples at 1, 7, and 14 days after introduction. Bt expression profile from D1 clustered distinctly from that of D7 or D14 (p-value < 0.05, pairwise PERMANOVA), whereas D7 and D14 profiles were statistically indistinguishable.

First, we addressed the question of whether there are differences in *Bt* gene fitness at different times after introduction into the mouse gut. We performed Principal Coordinates Analysis (PCoA) using Bray-Curtis dissimilarity on the relative abundance of the RB-Tn mutant strains within each mouse across the days of the functional genetics experiment and found that across multiple independent cohorts of this experiment, the mutant pool composition shifted across time in a replicable, largely deterministic pattern (Fig. 1B). PERMANOVA analysis showed that the data cluster significantly by experimental day (p = 1e^-4^, R^2^ = 0.457). Qualitatively, the largest shifts in PC1 occurred between D1 and D7, whereas the mutant pool changed less dramatically between D7 and D14. Consistent with the global changes in mutant abundance with time, clustering of WT *Bt* gene expression data from our transcriptomics experiment reveals that the largest changes in expression occur during the first week after introduction of *Bt* (Fig. 1C). The *Bt* expression profile from D1 clustered distinctly from that of D7 or D14 (p-value < 0.05), whereas D7 and D14 profiles were statistically indistinguishable. Together, these data suggest that a large shift occurs early during the first week of colonization such that different sets of genes mediate colonization and growth of the organism before and after the switch.

### Amino acid, vitamin, and capsular polysaccharide biosynthesis are transcriptionally upregulated and functionally significant during early invasion of the gut

To determine the scale of the observed shifts in gene usage, we compared the transcriptomes of WT *Bt* isolated on D1 and D7. Differentially expressed genes (DEGs) were identified for each pairwise comparison using the parameters logFDR < -3, |log_2_FC| > 2, and max group mean > 50 TPM. Even using these stringent criteria, we identified a staggering number of DEGs. 154 genes were significantly enriched on D1 compared to D7, while 359 genes were significantly enriched on D7 compared to D1 (Fig. 2A).

**Figure 2:**
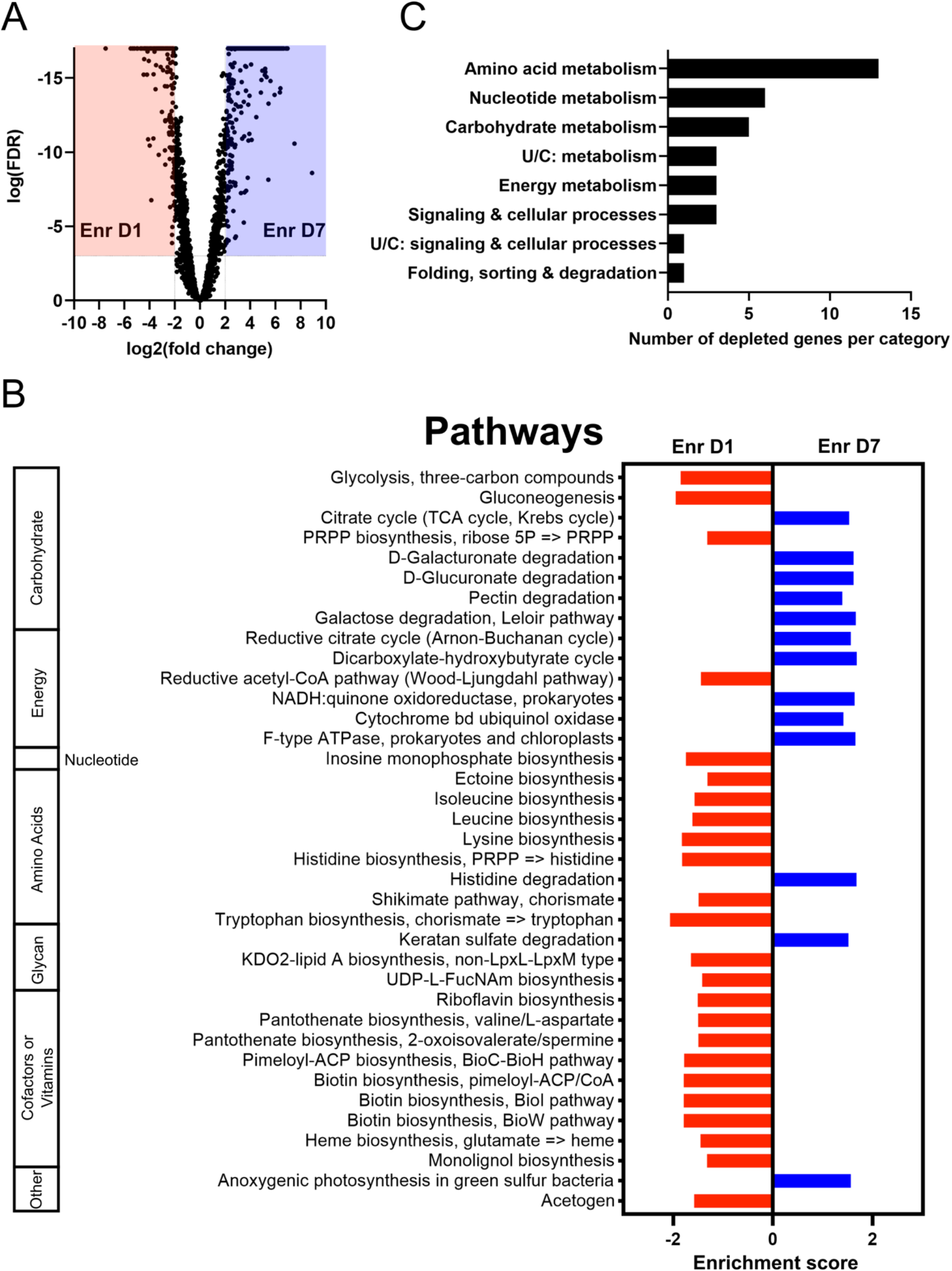
The Bt transcriptome undergoes dramatic remodeling during the first week after introduction to the murine gut. **A)** Volcano plot of significant transcriptional differences between D1 and D7 in WT Bt. There were 154 genes significantly enriched on D1 compared to D7 (red), while 359 were significantly enriched on D7 compared to D1 (blue) [cut-off: log(FDR-adjusted p-value) < -3, |log2(fold change)| > 2, and max group mean > 50 TPM]. B) Number of genes mutants significantly depleted in the RB-Tn experiment (t-statistic < -3σ) on D1 of the experiment. Each gene was assigned a KEGG functional category. Only genes with negative fitness are plotted because there were no genes with a positive fitness score / t-statistic on D1. C) Gene set enrichment analysis (GSEA) of all metabolic pathways in the Kyoto Encyclopedia of Genes and Genomes (KEGG) catalog, comparing D1 to D7 expression. Only pathways with significant differential expression between D1 and D7 (p<0.05) are shown.

Next, we asked whether the genes with higher relative expression on D1 correspond to specific functions. Though the period immediately following invasion of an exogenous organism has been studied in pathogenic infection contexts (13–15), very early time points in commensal colonization have been neglected in existing studies. This is a critical time for the invading organism, in which it must quickly adapt to a different set of stressors unique to the host environment, including differences in nutrient availability, pH, salinity, and drivers of the host immune system (16).

Previous attempts to characterize *Bt* colonization within the gut suggest that remodeling of the outer membrane is one response to the broad array of new selective pressures in the extracellular environment as the cell enters a new host (17–19). As is typical of other Bacteroides, *Bt* dedicates a substantial portion of its genome to its 8 distinct capsular polysaccharide (CPS) biosynthesis loci, which exhibit complex gene regulatory mechanisms. It has been shown previously that *Bt* can dynamically change its CPS expression profile *in vivo* (226), and that CPS4 plays a critical role in murine gut colonization (18, 19). Indeed, we saw major differences in expression of CPS biosynthesis genes between D1 and D7 (Fig. S1A). CPS1, 3, 4, and 8 loci have higher relative expression during the first day following introduction of *Bt*, whereas CPS5 and 6 are significantly enriched at D7 and beyond (Fig. S1B). Interestingly, mutants with disruptions in CPS1, 3, and 8 loci were not significantly depleted during the early days of the functional genetics assay. Only CPS4 mutants showed severe fitness defects (Fig. S2). This could indicate that CPS1, 3, and 8 serve redundant functions, or that fitness corresponds to protein levels or activity that are regulated independently of mRNA levels. Nevertheless, these results suggest that regulation of CPS gene expression is a response to the environmental changes that *Bt* encounters upon initial invasion of the murine gut.

To gain a more comprehensive understanding of the gene pathways expressed during the acute phase of adaptation to the host gut, we mapped the genes in the *Bt* genome to the 181 metabolic pathways within the Kyoto Encyclopedia of Genes and Genomes (KEGG) catalog and performed Gene Set Enrichment Analysis (GSEA) for each pathway on the differences in transcript abundances between D1 and D7 (Fig. 2B). The global expression pattern of *Bt* during D1 is reminiscent of the stringent response, in which growth is inhibited under conditions of nutrient limitation in favor of amino acid biosynthesis.

Analysis of gene expression changes reveals that pathways corresponding to the biosynthesis of many essential amino acids were enriched specifically at the D1 timepoint, including those for histidine, lysine, leucine, and isoleucine. This aligns with the findings of Watson *et al.* (20), which identify biosynthesis of essential amino acids as one of the main drivers of microbial colonization outcomes after fecal microbial transplant. Concurrently enriched at D1 were genes involved in the biosynthesis of biotin, which is a cofactor required for many reactions, including amino acid biosynthesis. These results are supported by RB-TnSeq analysis: by mapping the significantly depleted mutants (t-statistic < -3σ) on the first day of the functional genetics assay using KEGG Orthology (KO), we identified amino acid metabolism as the KO group with the largest number of significantly depleted mutants (Fig. 2C). These genes are identified by gold stars on the amino acid biosynthesis pathway map in Fig. 3A, which illustrates the individual reactions whose gene expression is enriched at D1 compared to D7. We observe that biosynthesis of most amino acids have either multiple reactions in the pathway that are transcriptionally enriched at D1, or one step that is functionally essential. Not only do both our transcriptomics analysis and functional genetics screen support a key role for amino acid biosynthesis early in colonization, but metabolomic analysis of the cecal contents corroborates this finding as well. We measured the levels of specific amino acids in the murine cecum before, post-D1, and post-D7 colonization, and found that amino acid levels were generally much higher D1 after colonization than D7 or D14 (Fig. 3B). This difference was especially profound and statistically significant for amino acids in the glycine-serine-threonine pathway, as well as aspartate. By contrast, glutamate and proline accumulate within the cecum during the first week of the time course before being depleted in the second week. The biosynthesis of glycine, glutamate, and aspartate was found to be functionally significant to the survival of *Bt* in the acute phase of colonization (Fig. 3A).

**Figure 3:**
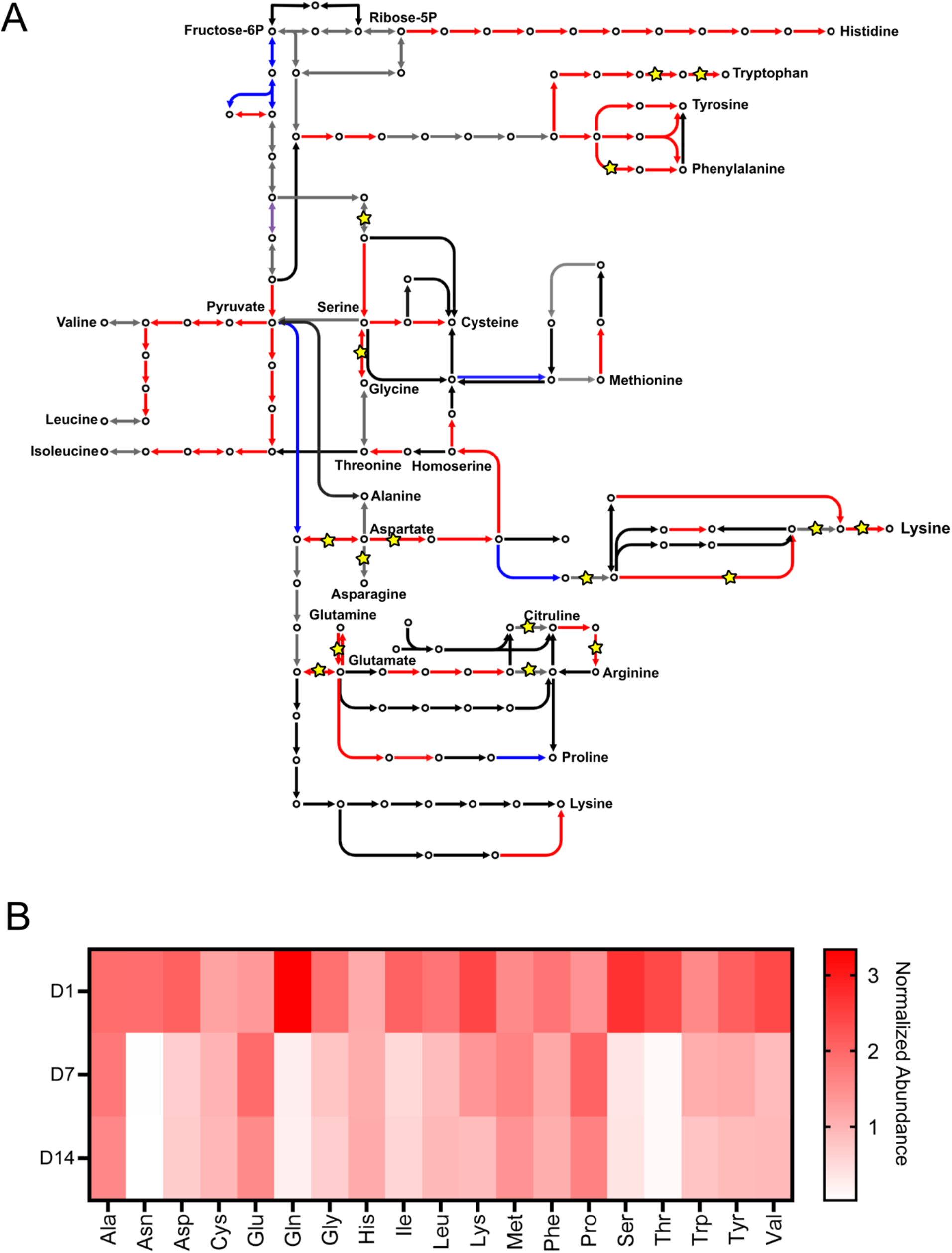
Biosynthesis of amino acids is transcriptionally enriched and functionally significant to early invasion success. A) Pathway map of reactions related to amino acid biosynthesis. Red arrows represent genes whose transcript is significantly enriched on D1, while blue arrows represent genes whose transcript is significantly enriched on D7 (FDR-adjusted p-value < 0.05). Genes whose transcript level did not differ significantly between D1 and D7 are colored in grey. Black arrows represent reactions for which the associated gene is unknown in the Bt genome. Gold stars represent gene disruptions that were depleted in the RB-Tn assay, as in Fig. 2B. B) Abundance of various amino acids in the ceca of germ-free mice measured using GCMS and normalized to internal standards and GF D0 controls (n=5 mice for GF D0, n=4 for post-colonization samples).

Interestingly, while amino acid biosynthesis was the predominant functional hit among significantly depleted mutants at D1 in the functional genetics assay, over the next two days, amino acid biosynthesis becomes less essential to cell fitness as *Bt* begins to resume its carbohydrate metabolism activities around D3, after overcoming an initial period of apparent starvation (Fig. 2C, Fig. S3).

### A shift toward enhanced expression of diverse sugar metabolism genes occurs during the first week of gut colonization

Members of the genus *Bacteroides* are well-known for their ability to digest a wide variety of polysaccharides. According to the CAZy database, *Bt* possesses 359 glycoside hydrolases, 87 glycosyl transferases, 15 polysaccharide lyases, and 19 carbohydrate esterases. Among the DEGs enriched on D7 compared to D1, carbohydrate metabolism stood out as the most prominently enriched functional category (Fig. S4). After the acute phase of colonization, *Bt* shifts toward expressing a remarkable number of polysaccharide utilization loci (PULs). Of the 42 total PULs that were significantly enriched at any point of the experiment, 34 were enriched on Day 7 or 14. These upregulated PULs encode genes belonging to a variety of pathways, including glycolysis, galactose degradation, pectin degradation, and D-galacturonate degradation (Fig. 4). Included among those are also PULs known to be involved in degradation of O-glycans (21), which suggests that *Bt* is foraging for sugars through digestion of the mucus layer. These results, along with a 10-fold increase in CFU in the latter half of the experiment (Fig. S5), suggest that within a week after introduction into GF mice, *Bt* recenters its transcriptome from a stringent response-like expression profile to a response centered predominantly on metabolism of the available sugars within the host gut.

**Figure 4:**
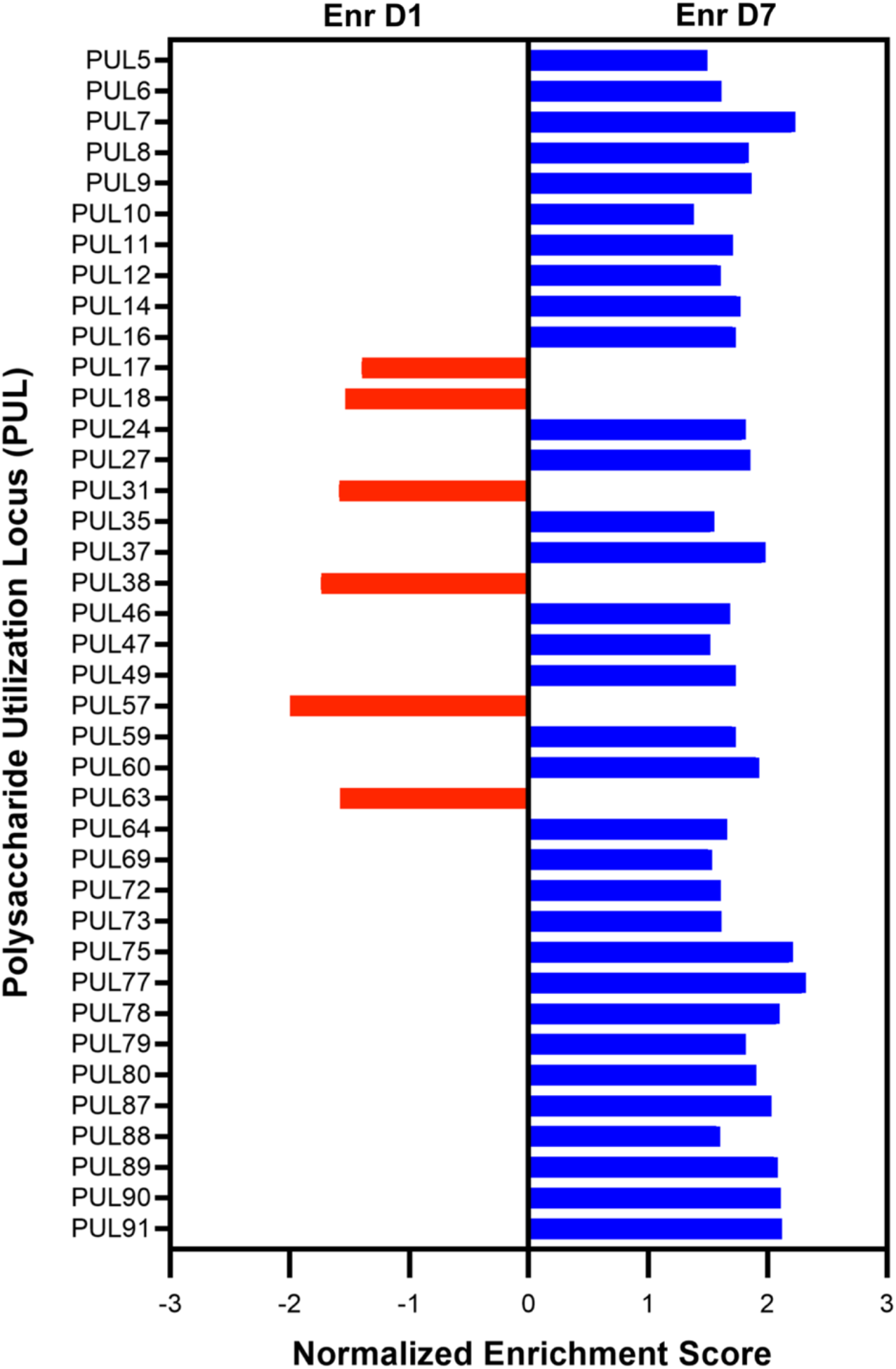
A shift toward higher relative expression of diverse sugar metabolism genes occurs during the first week of gut colonization. GSEA for genes in the 88 polysaccharide utilization loci of Bt. Only the significant PULs are shown. Of the 42 total PULs that were significantly enriched at any point of the experiment, 34 were enriched on Day 7 or 14.

### Upregulation of α-galactosidase activity confers a significant growth advantage to Bt in GF mice fed a standard RFO-rich diet

In stark contrast to the large transcriptional shifts that occurred in the first week after introduction, the transcriptional profile at D7 is mostly sustained through the rest of the experiment (Fig. 5A). In other words, the transcriptome of host associated *Bt* has largely stabilized by the end of the first week. The one salient exception in the second week is the continued upregulation of PUL24, suggesting that PUL24 may enable *Bt* to gain a competitive growth advantage as it proceeds to the persistence stage of engraftment. PUL24 has an operon structure typical of a PUL, with regulatory elements, transporter proteins and carbohydrate-active enzymes (Fig. 5B). Notably, the last gene of PUL24 encodes an α-galactosidase (BT1871), which is predicted to confer the ability to hydrolyze the α-1,6 glycosidic linkage in raffinose family oligosaccharides (RFOs), a major component of the fiber-rich diet that our mice were fed (22). Though this study focuses on understanding the shifting genetic determinants of colonization over time, we recognize that the effect of diet is intrinsic to the results of any gut microbiome study.

**Figure 5:**
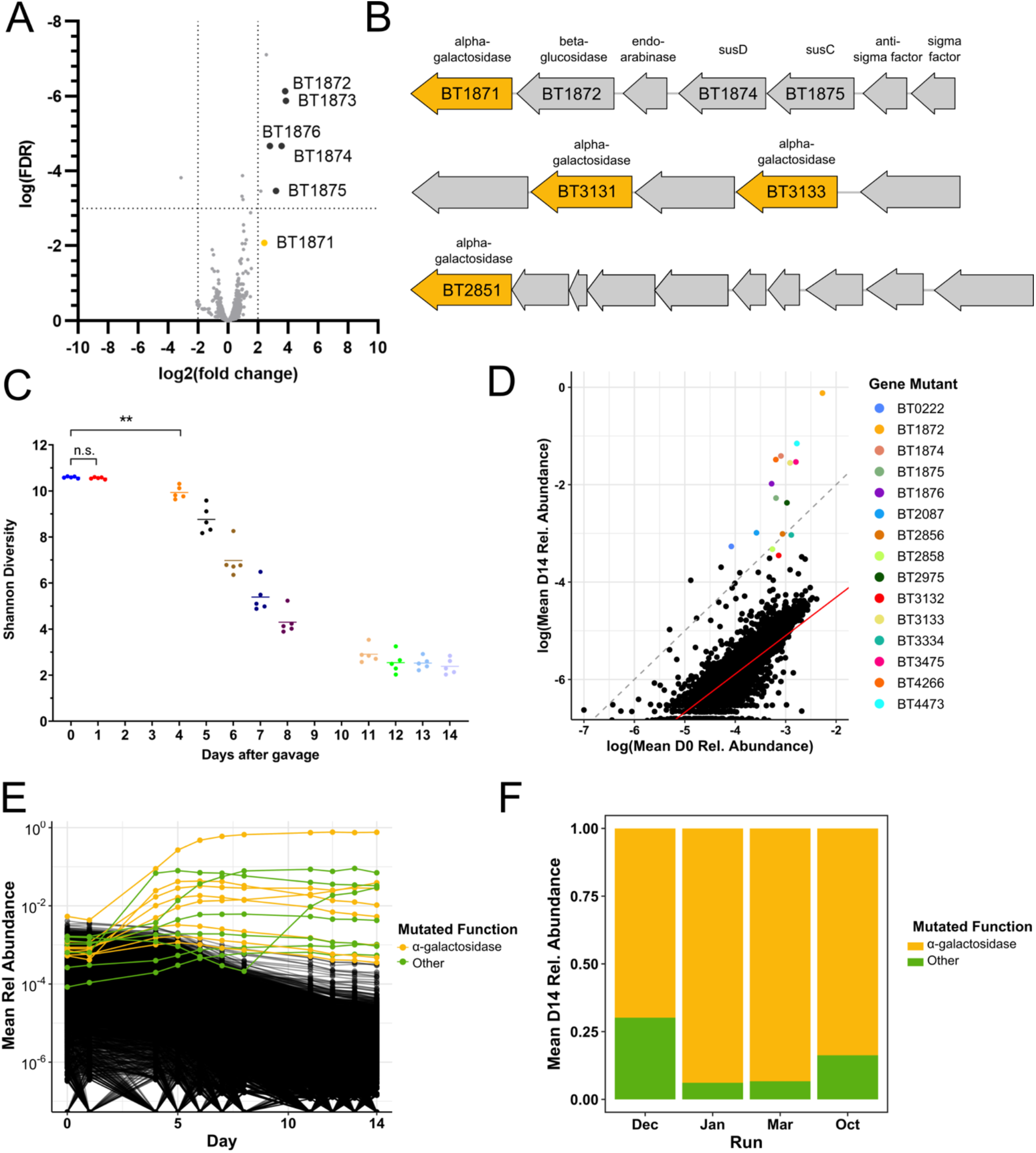
Colonization of a complex Bt mutant library within GF mice selects for disruptions upstream of alpha-galactosidase genes. A) Volcano plot of significant transcriptional differences between D7 and D14 in WT Bt. Using the same cutoff criteria as in Figure 2A, expression of all genes in PUL24 (red points) except for BT1877 increased significantly throughout the two weeks of the experiment. B) Organization of PUL 24 (BT1871-1877), PUL 39 (BT2851-2860), and an unnamed PUL containing BT3130-3134. Predicted alpha-galactosidases are colored dark yellow. C) Shannon diversity of the RB-Tn mutant pool in mice over time. Each point represents the RB-Tn pool within a single mouse on a particular day. The mice are initially colonized by an extremely diverse pool of mutants (300,000+ strains) but diversity drops significantly by D4 (mean = 114,256 strains) and continues to fall throughout the experiment (mean = 20,269 strains by D14). D) Initial (inoculum) versus final (D14) relative abundance of RB-Tn mutant strains for a single experimental run (October). Each point represents the summed relative abundance of all mutant strains that corresponded to a given gene, averaged across mice. Dashed grey line designates a 1:1 relationship between starting and final abundance; red line represents a linear regression best-fit line generated from the log-transformed data (p < 2e-16, R^2^ = 0.6461, Methods). E) Relative temporal abundance of RB-Tn mutant strains from a single experimental run (October). Each line represents the summed relative abundance of all mutant strains that mapped to a given gene, averaged across mice. Top 15 most abundant gene mutants at D14 are colored; all others are black. F) D14 fractional abundance of gene mutants mapping to operons that encode alpha-galactosidase functions (yellow) or other gene functions (green) averaged across mice for each experimental run.

Our functional genetics screen confirmed the importance of PUL24: within the first five days after introduction of the RB-Tn mutant library into GF mice, the diversity of the community quickly collapses (Fig. 5C) due to strong positive selection for a small pool of hyperfit mutants. RB-Tn mutants with insertions in PUL24 were among the most enriched mutants at the end of the two-week experiment, even after controlling for their relatively high abundances in the inoculum (Fig. 5D). Furthermore, most of the significantly enriched mutants had Tn insertions in one of three operons: PUL24 (BT1871-1877), PUL39 (BT2851-2860), or BT3130-3134, all of which encode at least one α-galactosidase, situated at the tail end of the operon (Fig. 5B). At the end of the Tn selection experiments, the populations were overtaken by mutants carrying Tn insertions upstream of these α-galactosidase genes (Fig 5E). This was true for all four independent experimental runs, which were performed months apart from one another (Fig 5F).

We note that *in vitro* passaging of this RB-Tn mutant library within BHI-S medium also resulted in a modest but significant decrease in Shannon diversity over two weeks; however, the mutant library was able to maintain alpha diversity *in vitro* when passaged in the presence of a specific-pathogen-free (SPF) community (Fig. S6). Engraftment of the RB-Tn mutant library in SPF mice was limited by a severe bottleneck, but the rare population that successfully penetrated the gut was stably maintained at low abundance throughout the course of the next two weeks (Fig. S7). These patterns suggest that gene-environment interactions for *Bt* are affected by the presence or absence of other community members.

This RB-Tn pool has previously been assayed in over 300 different conditions including distinct carbon or nitrogen sources and specific stress conditions (16). The mutants that we identified as being hyperfit in GF mice exhibit a phenotype significantly deviant from WT in only two conditions: within GF mice and in defined culture with melibiose—a disaccharide of glucose and galactose—as the sole carbon source. In both these cases, these mutants exhibit a growth advantage. Indeed, when we isolated the most abundant strains from D14 of the RB-Tn experiment, we found that their growth rate was significantly faster than WT when melibiose is the sole carbon source (Fig. 6A). In this condition, *Bt* must hydrolyze the α-1,6 glycosidic bond to harvest and metabolize the monosaccharide sugars. The competitive advantage of these mutants cannot be attributed to either of the monosaccharides, as the mutant growth rates with these carbon substrates are indistinguishable from that of WT (Fig. S8).

**Figure 6:**
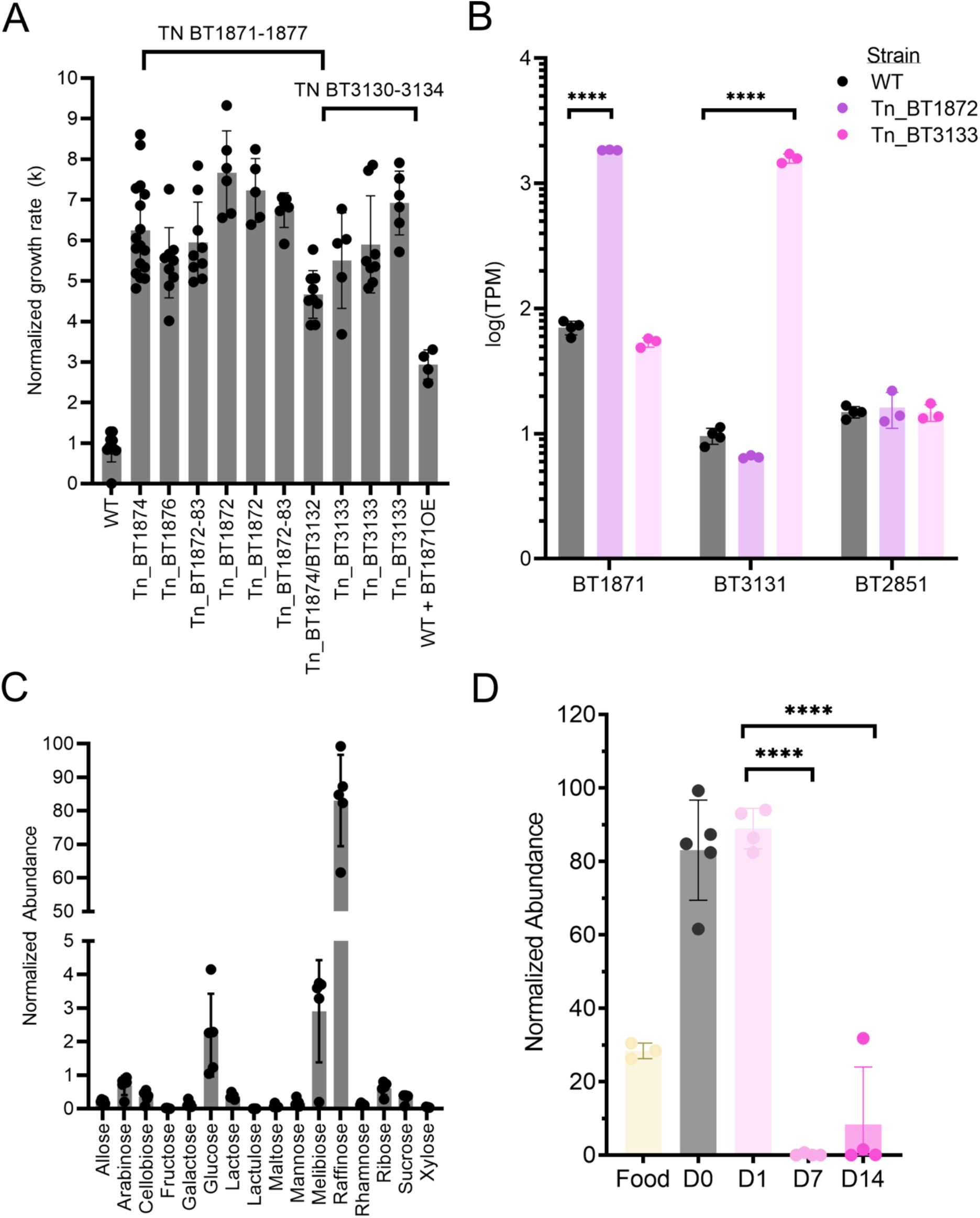
Upregulation of α-galactosidase activity confers a significant growth advantage to Bt in GF mice fed a standard RFO-rich diet. A) Log phase grow rate (k) of the most abundant RB-Tn mutant strains isolated from mice. Strains with asterisk carry other mutations in addition to the transposon insertion. The WT + BT1871OE strain carries a plasmid copy of BT1871 expressed from the *rpoD* promoter that was integrated into the WT genome at the attN1 site. B) Normalized abundance of alpha-galactosidase mRNA (transcripts per million, TPM) were measured in WT and hyperfit RB-Tn mutants isolated from mice and grown in Varyl-Bryant defined medium with 20mM melibiose as the sole carbon substrate. Expression of the alpha-galactosidase BT1871 was more than 10 times greater with an upstream insertion in BT1872 (Tn_BT1872) compared that of WT, and expression of the alpha-galactosidase BT3131 was more than 100 times greater with an upstream insertion in BT3133 (Tn_BT3133) compared to WT. Expression of alpha-galactosidase BT2851 was unaffected in either of the mutant strains, since the transposon insertions were not upstream of this gene. C) Abundance of various sugars in the GF mouse ceca as measured using GCMS and normalized to internal standards. D) Abundance of raffinose in the standard chow fed to GF mice, within GF ceca before colonization, or 1, 7, or 14 days post colonization.

Given that release of the monosaccharide sugars of melibiose depends on hydrolysis of an α-1,6 glycosidic bond, we wondered if the fitness phenotypes in both mice and *in vitro* depended on overexpression of the α-galactosidase gene downstream of the Tn insertion site. The original publication characterizing this *Bt* RB-Tn library (23) showed that polar effects can be expected in which readthrough from the strong promoter driving antibiotic resistance in the transposon can overexpress genes downstream from the insertion. To test the hypothesis that the transposon insertion enhanced expression of downstream genes, we grew WT and hyperfit mutants in melibiose medium and measured the expression of α-galactosidase genes. For this experiment, we used two mutants, one carrying an insertion in PUL24 and another carrying an insertion in the BT3130—3134 operon. We found that, in both cases, expression of the α-galactosidase downstream of the insertion but within the same operon was overexpressed by at least 10 times compared to WT (Fig. 6B). Meanwhile, expression of an α-galactosidase (BT2851) located outside the operons carrying the insertion was similar between WT and the mutants. We then overexpressed *BT*1871 from a strong, constitutively active promoter (P_rpoD_) in WT *Bt*, and observed a three-fold increase in log-phase growth rate when melibiose was the sole carbon source (Fig. 6A).

Metabolomic measurements of a carbohydrate panel confirms that high concentrations of raffinose and its constituent sugars are among the most abundant substrates within the ceca of GF mice fed a standard chow (Fig. 6C). Raffinose and melibiose build up in the lower GI tract of these mice at high concentrations until *Bt* is introduced. After 7 days, during which time *Bt* initiates overexpression of α-galactosidase genes, these sugars are depleted (Fig. 6C, Fig. S9A). In contrast, the other disaccharide product of raffinose, sucrose, which does not have an α-1,6 glycosidic bond, is consumed by the host and no significant changes are observed in sucrose concentration following the *Bt* colonization (Fig. S9B).

Together, we find that when mice are fed a standard high-fiber chow, the high concentration of RFOs accumulating in the lower GI tract creates an environment that strongly selects for *Bt* strains that can make efficient use of these sugars through increased expression of α-galactosidases. We infer that efficient carbohydrate metabolism—particularly for abundant dietary fibers—is a major determinant of population-level selective dynamics during the persistence phase of *Bt* engraftment within the gut.

### Strong selection for efficient RFO metabolism leads to emergence of spontaneous Bt mutants with duplicates of the BT1871 locus through an IS3-family transposable element

The transposon mutants in our functional genetics experiment were able to gain ∼10-fold increase in α-galactosidase expression due to readthrough from an extremely strong, synthetic promoter. We wondered if the selective pressure for elevated α-galactosidase activity to utilize the α-1,6-linked sugars abundant in the diet would drive evolution of a spontaneous mutant with enhanced α-galactosidase activity. We inoculated three GF mice with WT *Bt*. After one week, they were each separated into individual cages. At the end of the six-week incubation period, we performed shotgun metagenomic sequencing on fecal material from all three mice. Assembly of the short-read sequences revealed that there was at least 2x coverage of the BT1871 locus for all samples (Fig. 7A). A dip to 1x coverage in the middle of this region mapped to an IS3-family transposase, of which there are six other identical copies in the genome; hence, the corresponding coverage was diluted among the different copies. BreSeq analysis (24) identified three new junctions in the *Bt* genome indicative of genome rearrangements (Fig. 7B). All three junctions were formed between either BT1872 or BT1873 and a locus downstream of BT1871, which encodes for rRNAs, tRNAs, and a ribosome recycling factor. All three new junctions occupied a significant portion of the sequencing reads (60-73%), suggesting that the mutants carrying these junctions were sufficiently competitive in the gut environment to comprise a significant portion of the fecal *Bt* population.

**Figure 7:**
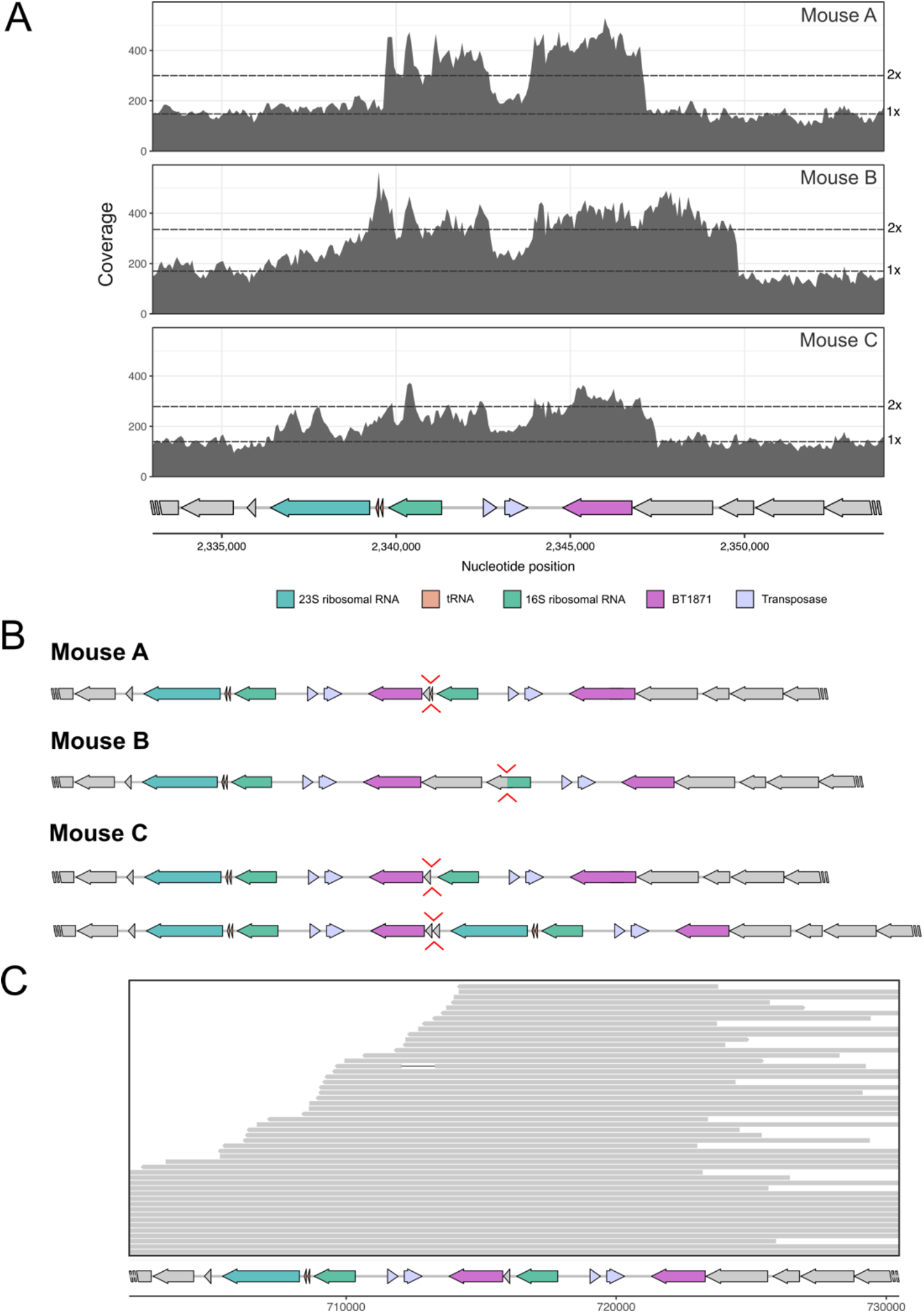
Under strong selective pressures, Bt duplicates the BT1871 locus with the help of a transposable element. A) Shotgun whole genome sequencing of bulk fecal samples from three mice inoculated with WT Bt for 6 weeks shows at least 2x coverage of the BT1871 locus. B) For each mouse, Breseq analysis of the shotgun whole genome sequencing identified a new junction joining a segment up- and downstream of BT1871 in 60-70% of the reads. C) Long-read MinION sequencing of an abundant mutant in Mouse C (MZ65) reveals the duplication of a long segment including the BT1871 gene, which generates a tandem repeat of the locus. Gray bars represent individual long reads, which together span the region containing the tandem repeat.

IS3-family transposases are known to act by a copy-and-paste mechanism (25). We wondered if this transposase could have created multiple copies of the BT1871 locus. We isolated individual mutants from the bulk fecal material, cultured the isolates in complex broth (BHI-S), and performed long-read sequencing with MinION. In one of the isolates, we identified a tandem repeat of the BT1871 locus (Fig. 7C), which places the original copy of BT1871 downstream of a strong rRNA promoter. Like the RB-Tn mutants, the mutants isolated in this experiment all grew about five times faster than WT in defined minimal media with melibiose (Fig. S10). Interestingly, the six identical copies of this IS3-family transposase are often found adjacent to PULs in the WT *Bt* genome. This suggests that these transposable elements confer a mechanism to modulate gene expression that may be advantageous in adapting to new metabolic landscapes.

## Discussion

### Microbial adaptation during colonization is a dynamic process

Bacteroides species are dominant and prevalent in many human populations. Instrumental to their success is their ability not only to tolerate stress, but to quickly adapt in order to grow under a range of conditions. *Bt* has been used as a model organism to understand the genetic drivers of gut colonization for commensal organisms (19, 26, 27). However, by and large, a single representative timepoint has been assessed in previous investigations, which fail to take into account the dynamism of the adaptational process. For example, contrary to the established characterization of *Bt* as exhibiting constant, rapid growth in the gut based on gene expression at a single timepoint (28), our time course data evaluating CFU counts (Fig. S5), transcriptomics, and functional genetics suggest that upon entry into the gut ecosystem, *Bt* growth lags for at least 4 days, during which *Bt* prioritizes biosynthesis of amino acids and other essential compounds, before it begins a period of rapid growth. In this study, we find that the set of genes required for *Bt* to initially establish colonization in the gut are distinct from those required to persist (Fig. 1). Though the findings of previous studies do reflect a replicable snapshot of the *Bt* adaptational process, these studies have largely neglected the ecological dynamics of *Bt* adaptation that play out temporally on the scale of weeks. The results presented here suggest that because of the rapidly shifting adaptational profile exhibited by microbial populations over the course of colonization, investigators must take care in selecting the appropriate timepoints to address their experimental questions.

### Dynamically changing pathways in Bt

The process of colonizing a new host presents a diverse array of environmental changes that require physiological adaptation. For example, the bacteria experience changes in pH and nutrient availability as well as assaults by the host immune system. The largest transcriptomic changes that we observed across the two weeks of *Bt* colonization occurred in the first week: 10% of all *Bt* genes (503 genes) were differentially expressed between D1 and D7, whereas only 0.1% of the genes were differentially expressed between D7 and D14 (Fig. 2A).

Our findings suggest *Bt* faces nutrient starvation upon arrival in the lower GI tract during the earliest stage of colonization. Under these conditions, microbes activate a program of transcriptional changes called the stringent response, in which carbohydrate metabolism is curtailed in favor of amino acid biosynthesis. Schofield *et al*. (29) previously showed that *Bt* requires the stringent response to establish and maintain colonization in ex-germ-free mice. Indeed, in this study we demonstrated that biosynthesis of amino acids is both highly upregulated and functionally vital to the initial establishment of *Bt* colonization. One possible explanation for why amino acid biosynthesis pathways have significantly higher expression on D1 compared to D7 or D14 is that when *Bt* first arrives in the uncolonized gut, dietary or host-derived amino acid resources are scarce enough to induce the stringent response, which leads to the general upregulation of amino acids. Upon adjustment to the environment, it will be more energetically favorable to fine-tune expression of only the necessary resources.

In addition to amino acid biosynthesis, the other major category of genes that were highly expressed on D1 but not D7 or D14 are those involved in vitamin biosynthesis. Genes including those related to biotin (Vitamin B7), riboflavin (Vitamin B2), and pantothenic acid (Vitamin B5) biosynthesis were significantly enriched on D1. These vitamins may simply be scarce in the lower GI tract early during colonization of ex-germ-free mice, requiring *Bt* to synthesize them itself early in colonization. Indeed, dietary B vitamins are primarily absorbed by the host in the small intestine, and although microbe-derived B vitamins can be produced and absorbed in the colon, in our germ-free model, we would expect B vitamin levels to be low in the distal GI tract prior to colonization (30). It is also possible that the upregulation of biotin and other vitamins could be linked to host immunity. Given that microbial biotin biosynthesis genes are upregulated during the infection of many pathogens in humans, including pathogenic *E. coli* gastrointestinal infections and *Salmonella* bloodstream infections (31–33) and host biotin deficiency has been demonstrated to increase secretion of proinflammatory cytokines in human dendritic cells (34), upregulation of biotin biosynthesis by *Bt* may serve to dampen inflammation from the host in response to bacterial colonization. Though *Bt* is not traditionally regarded as a pathogen, we cannot rule out the possibility that it can induce some level of inflammation, especially in mice that have never encountered microbes. *Bacteroides fragilis* is a well-studied species of the same genus that is commonly part of a healthy microbiome but can induce inflammation in specific contexts (35, 36). Future investigation will be required to understand whether upregulation of vitamins in *Bt* is related to any of these host responses.

### Loss of diversity in RB-Tn pools *in vivo* and *in vitro* and the impact of complex communities

In both the mouse gut and in complex broth (BHI-S), our RB-Tn mutant pool exhibited significant losses in Shannon diversity during the first week, which extended through the remainder of the two-week experimental period (Fig. 5C; Fig. S6). Whereas *in vitro* loss of diversity was primarily attributable to shifts in the relative abundance of the mutants rather than extinction of mutant strains, the collapse in diversity observed in the host-associated RB-Tn pool was caused by decreases in both strain evenness and richness— and on a much larger scale than in the *in vitro* experiments. This difference in scale suggests that *in vivo,* strong, directed selective pressure for improved melibiose and RFO metabolism led to population sweeps by strains carrying gain-of-function mutations. By contrast, the loss of diversity exhibited by our *in vitro* experiments suggests a much weaker and more diffuse selective environment, which is not surprising given that the mutant population was generated and has been cultivated *in vitro* in BHI-S. Moreover, WT *Bt* already grows efficiently in BHI-S; thus there is limited dynamic range to respond to the selective pressures of that environment with improved growth. By contrast, WT *Bt* growth on melibiose is highly inefficient, with a doubling time of around 10 hours, and the RB-Tn gain-of-function mutations were able to increase growth speed on melibiose by log-fold differences.

A recently published transposon screen for enteric colonization factors in *Klebsiella pneumoniae* similarly reports that despite robust colonization of the colon by a diverse array of mutants at D1, the diversity of the mutant population was greatly reduced by D4 and there was marked expansion of a small subset of mutants (37). However, contrary to our findings, diversity of the mutant *Klebsiella* population recovered shortly thereafter. This difference can potentially be accounted for by the fact that the Klebsiella experiments were carried out in SPF mice treated with antibiotics that reduced, but did not eradicate, the native microbiome to allow for dense *Klebsiella* colonization.

We find that, *in vitro*, addition of an SPF community prevented the loss of diversity while passaging the *Bt* RB-Tn library for 14 days (Fig. 6). In accordance with these observations, a recent publication investigating the impact of community complexity on evolution of focal strains concluded that microbes have a higher evolutionary capacity in low diversity communities than in complex communities (38). Thus, the strong, directed selection observed in our *in vivo* experiments may be rendered more diffuse in the presence of a native microbial community.

Among other factors, we expect that the results of this study will likely differ depending on the presence or absence of a pre-established community. Though we recognize the limitations of studying adaptation and colonization in GF mice, these experiments lay foundational groundwork for the eventual investigation of community assembly and fitness dynamics in more complex communities. Specifically, having a thorough catalog of host-specific colonization dynamics in the GF gut will create a basis of comparison from which we can delineate host-versus microbiome-induced selective pressures and adaptive dynamics. Preliminary assessments of RB-Tn mutant engraftment in complex communities revealed that a severe initial bottleneck in colonization led to very low levels of *Bt* persistence in the community (Fig. S5). Moreover, the final composition of mutant strains appeared to be dictated by stochastic forces, with high inter-individual variability, and no consistent selective pattern for particular genes or functions (Fig. S7). This further supports the notion that addition of a complex community may limit the extent of strong, directed selection, although additional follow-up experiments are required to further characterize these patterns.

### IS3 transposable elements: a potential novel mechanism to modulate expression of specific CAZymes?

In GF mice fed a standard RFO-rich diet, the selective pressure for mutants with increased α-galactosidase activity is severe. Mutants in the BT1871 locus were consistently selected for in WT *Bt* allowed to evolve in mice for six weeks. In all three populations, the downstream side of the duplicated region ends midway through a ribosomal gene, meaning that there are no known transcriptional terminators between the ribosomal promoter and the α-galactosidase gene, BT1871. It is likely that these mutants have increased α-galactosidase activity due to readthrough from the strong ribosomal promoter in the upstream copy of the locus. Though increased α-galactosidase could be simply due to the presence of two copies of BT1871, the growth phenotype that these mutants exhibit, in which log phase growth is 5x faster than WT, is on par with that of the RB-Tn mutants, for which α-galactosidase expression was more than 10 times greater than that of WT.

In *Bt,* identical copies of this IS3 transposable element occur in seven locations, often adjacent to PULs and almost always paired with a ribosomal gene. It is well known in both mammals and prokaryotes that transposable elements can modulate the expression of nearby and distant genes. Transposable elements may be selected for because the genetic “cargo” that they shuttle along (such as an antibiotic cassette) is beneficial to the organism or because the position where the insertion occurs results in some downstream effect that is beneficial to fitness. Both reasons may contribute to the selection of the tandem duplication of the BT1871 locus. Future studies will be necessary to investigate whether other IS3 elements in *Bt* perform similar functions in modulating expression of nearby genes and whether such a mechanism can be found in other families of bacteria.

## Materials and Methods

### Microbial Strains and Growth Conditions

The bacterial strains used in this study are listed in Table S1 in the supplemental materials. *Bacteroides thetaiotaomicron* VPI-5482 [AMD595, (23)] and all its derivatives were cultured anaerobically at 37°C in liquid BHI-S medium or defined Varyl-Bryant medium as described in Ref 23. Varyl-Bryant medium with no carbon source was supplemented with glucose, galactose, raffinose, or melibiose to a final concentration of 20mM. An anaerobic chamber (Coy Laboratory Products) containing 20% CO2, 10% H2, and 70% N2 was used for all anaerobic microbiology procedures.

### Mice

Female mice 8-12 wk-old C57Bl/6J GF mice were bred and maintained in plastic gnotobiotic isolators or bioexclusion racks within the University of Chicago Gnotobiotic Core Facility and fed ad libitum autoclaved standard chow diet (LabDiets 5K67). The mice were given a single dose of either wildtype *Bt* VPI-5482 or the *Bt* mutant library (gift from Deutschbauer lab) at 106-108 CFU / 200μL. All murine experimental procedures were institutionally approved.

### *In vivo* genome-wide mutant fitness assays

Several individual cohorts of mice were used, indicated by the month of the experiment. For each cohort, mice were housed in a single gnotobiotic isolator for the duration of the experiment (December and January), or each individual cage was housed in a bioexclusion rack (March and October). Female C57BL/6J GF mice between 8-12 weeks old were fed a standard irradiated diet ad libitum, split into cages of n = 2-3 mice/cage, and allowed to acclimate for 3 days prior to colonization. The inoculum was prepared by one of two methods: 1) for Mar. and Oct. experiments: thawing a 2mL aliquot of the *Bt* RB-TnSeq library, growing the entire aliquot in 150mL BHI-S medium overnight with 20 ug/mL erythromycin (16h), and backdiluting the culture to OD600=0.05 the next morning to allow for cells to reach mid-log phase (3hr), or 2) for Dec. and Jan. experiments: thawing aliquots of the *Bt* RB-TnSeq library and gavaging the thawed *Bt* cells directly into mice. For each experiment, at least 3 cell pellets of the inoculum were collected at Time=0 references. Each mouse was colonized by oral gavage with 200μL of the *Bt* transposon library. Stool samples were collected daily (excluding weekends), up to 14 days post-colonization to assess longitudinal shifts in mutant abundance. Mice were monitored and weighed daily. Genomic DNA was extracted using the DNeasy PowerSoil Kit.

### Pipeline for measuring relative abundance and fitness scores of RB-Tn mutants

RB-TnSeq strain and gene fitness scores were calculated as described previously from strain-level count data (39). For temporal abundance analyses of TnSeq mutants for each run, we first created a feature table with raw counts of strain-level mutants across each sample. This table was filtered to remove strains with counts of 1, as these are likely produced by sequencing error. Next, counts were normalized by the count of a synthetic spike-in barcode that was introduced at 20pM into each sample during PCR amplification of the barcodes. Samples where the spike-in represented > 30% of total reads were discarded. The synthetic spike-in barcode was subsequently removed as a feature from the table, and the resulting tables were used for alpha diversity analyses via the R package vegan. For all other relative abundance analyses, strains were assigned to genes based on previous mapping by Liu *et al*. (23), as well as manual mapping performed for this experiment. Strains that had not been previously mapped were binned together as “non-mapping” strains, and strain-level counts were then summed for each gene. For all subsequent analyses, we further filtered out genes that mapped to *Bt* plasmids, as we were more interested in chromosomal gene fitness patterns. The R package phyloseq was used to calculate Bray-Curtis dissimilarity for all pairwise combinations of samples, which was then used to create PCoA plots. Finally, filtered count tables were adjusted to relative abundance based on the total remaining counts, and used to track gene-level relative abundance over time. For linear regression of initial vs final relative abundance of gene mutants, genes with zero-counts were re-assigned a value of 1e^-7^ to perform log-transformation of the data.

### *In vivo* transcriptomic experiments

Mice were sacrificed at 1, 7, and 14 days post-colonization. Luminal contents of the mice cecum were immediately snap-frozen in liquid N2 and stored at -80C. About 50mg of the contents were transferred into 2mL screw-cap tubes, followed by the addition of 1mL TriZOL reagent for the isolation of RNA. The samples were homogenized by beadbeating with 0.1-mm glass beads in a Mini-BeadBeater-96 for 2 minutes. Total RNA isolation and purification were performed using the TRI reagent protocol and quality checked by BioAnalyzer. All library preparation and sequencing work was performed by MiGS (Pittsburgh, PA). Initial DNAse treatment is performed with Invitrogen DNAse (RNAse free). Library preparation is performed using Illumina’s Stranded Total RNA Prep Ligation with Ribo-Zero plus. Custom Ribo-Zero probes were designed for *Bt* and supplemented alongside the standard probe set. Custom probe sequences can be found in Table S2. Sequencing was performed on a NextSeq2000 giving 2x50bp reads. Post sequencing, we use bcl2fastq (v2.20.0.422) to demultiplex and trim adaptors.

### Gene Set Enrichment Analysis (GSEA) of metabolic pathways

Metabolism for the *Bt* genome was first estimated using the anvio-v7 program “*anvi-estimate-genome”* that identifies the KEGG Ortholog family (KOfam) annotations for each open reading frame. Gene calls for each metabolic pathway found within the *Bt* genome were then transferred into a unique GSEA pathway query list in R (*’fgsea’)*. Pathway enrichment was then calculated using the enriched gene lists derived from the RNA-Seq analyses using ‘*deseq2*’. Pathways were filtered for *padj>0.05* and the normalized enrichment scores (NES) plotted (PRISMv9). Other custom gene lists were created to calculate the enrichment of other gene sets including the polysaccharide utilization loci (PULs), genes involved the production of specific amino acids, and capsular polysaccharide loci (CPSs).

### Isolation of mutants from fecal matter

Mouse feces were collected and immediately homogenized in 500mL 25% glycerol solution and stored at -80C. Prior to isolation, glycerol stocks were allowed to thaw on the benchtop for 10 minutes, spun for 30 seconds at 2,000 RPM. On each 150mm BHI-S plate, 100uL of 10-3, 10-4, or 10-5 dilution of the glycerol stock was spread using 4.5 mm glass beads. The plates were incubated anaerobically at 37C for two days. Individual colonies were picked into 1mL 96-well plates containing 750uL BHI-S in each well.

After 16 hours of growth, glycerol was added to a final concentration of 20% and the isolates were stored. The isolate stocks were used as the template in PCR amplifying the barcoded region of the mutants. The PCR products were sent for Sanger sequencing.

### *In vitro* culture and growth measurements

*Bt* was grown in an anaerobic chamber at 37°C either in Brain Heart Infusion Supplemented (BHI-S) medium or Varel-Bryant (VB) defined medium. The Varel-Bryant medium base was made with no carbon source and was supplemented by 20mM of the indicated carbon source. For growth measurements, colonies of *Bt* were inoculated into 3mL of BHI-S in plastic culture tubes and grown overnight at 37°C, for a total of 6 biological replicates. Sealed Hungate tubes containing 10mL of VB medium and a 20mL headspace were used for subsequent growth. Immediately before inoculating with the cells, the tubes were inoculated via syringe with autoclaved sugar solution and hemin solution for a final concentration of 20mM sugar and 5 ug/mL hemin. Overnight cultures were diluted to OD600 = 1, and 100uL of the diluted culture was inoculated into the prepared media tubes via syringe. Anaerobically sealed cultures were grown outside of the chamber in a 37°C incubator with no shaking. Every 45 minutes, the cultures were taken from the incubator, cells resuspended by shaking, and their OD600 readings measured by a GENSYS 40-Vis spectrophotometer.

### Host-associated evolution of spontaneous Bt mutants

Three female mice 8-12 wk-old C57Bl/6J GF mice were co-housed in the same gnotobiotic isolator and fed standard chow diet ad libitum. One week post inoculation of *Bt*, the three mice were separated into individual cages. A fecal sample was taken six weeks post inoculation. DNA was extracted from the fecal pellets by the phenol-chloroform method, followed by ethanol precipitation, and sent for shotgun sequencing. Individual isolates from each fecal pellet were cultured from the bulk material as outlined in Isolation of mutants from fecal matter. We assayed for isolates with increased growth in VB-melibiose and selected one isolate from each mouse for MinION long-read sequencing.

### Isolated genomes sequencing, assembly and polishing

To provide greater context for the delineation of the complex chromosomal rearrangements associated with the BT1872/BT1873 operon, a long-read sequencing strategy was employed. The isolate genomes assessed were wild-type *Bt*, the strain used for the mouse experiments, and three spontaneous mutant cultivars (MZ55, MZ58 and MZ65), which demonstrated enhanced growth rates in the presence of melibiose, recovered from the feces of mice six weeks after initial inoculation. Total genomic HMW DNA was extracted by a standard phenol chloroform protocol (40) on overnight 25-mL BHIS broth cultures. DNA was resuspended in 0.1-mL 10 mM Tris-Cl, pH 8.5.

Slow pipetting, wide bore pipette tips and steps to minimize velocity gradients were implemented throughout to avoid further shearing of DNA molecules. Libraries were prepared with the Rapid Barcoding Kit (SQK-RBK004) and the standard protocols from Oxford Nanopore Technologies were used with the following modifications. DNA fragmentation was performed on 10-ug DNA using 10 passes through a 22G needle in a 250-*µ*L volume before purification using 0.5% Agencourt AMPure XP beads (A63882, Beckman Coulter). Each elution step of the AMPureXP beads was performed using 10 mM Tris-Cl pH 8.5 instead of water, at 37°C for 5 min. The gDNA inputs into library preparation ranged between 0.5 *µ*g and 1.2 *µ*g (Table S3), based on sample availability in a standard 8.5 *µ*l volume, with 1.5 *µ*l Fragmentation mix added to each sample. Barcoded libraries were pooled so each sample contributed an equal input mass (∼0.5 *µ*g, Table 9). Using MinKNOW (v4.3.4), a single R9.4/FLO-MIN106 flow cell (Oxford Nanopore Technologies) sequenced the final prepared library with a starting voltage of −180 mV and a run time of 72h. Guppy (v5.0.11) and the sup model were used for post-run basecalling, sample de-multiplexing and the conversion of raw FAST5 files to FASTQ files. For downstream analyses, we only used reads with a minimum quality score of 7. We assembled long-reads contigs with Flye (41). Additional DNA extractions were carried out for every isolate using a standard phenol-chloroform extraction and send for short-read sequencing. We then used the short-reads to polish the long-read assemblies using Pilon v1.23 (42).

### Metagenomic mapping and coverage visualization

We used anvi’o v7.1 (43) and the metagenomic workflow to compute and visualize metagenomics coverage for each isolate genome. Briefly, the workflow uses (1) Prodigal v2.6.3 (44) to identify open-reading frames (ORFs), (2) ‘anvi-run-hmm’ to identify single copy core genes from bacteria (n=71, (45) and ribosomal RNAs (n=12, modified from https://github.com/tseemann/barrnap) using HMMER v3.3. (46), (3) ‘anvi-run-ncbi-cogs’ and ‘anvi-run-kegg-kofams’ to annotate ORFs with the NCBI’s Clusters of Orthologous Groups (COGs) (47) and the KOfam HMM database of KEGG orthologs (KOs) (48, 49) respectively. We used Bowtie2 v2.3.5.1 (50) to recruit metagenomic short-reads to the contigs, and samtools v1.11 (51) to convert SAM files to BAM files. We profiled the resulting BAM files with ‘anvi-profile’ and used the program ‘anvi-merge’ to combine all single profiles into a merged profile for downstream visualization. We used ‘anvi-get-split-coverages’ and ‘anvi-script-visualize-split-coverages’ to generate the coverage plots. We used ‘anvi-export-gene-calls’ and gggenes v0.4.1 to visualize the genomics context around BT1871.

### BT1871 copy number

We used blast (52) to compute the number of long-reads with two copies of the BT1871 locus in the MZ65 isolate. We extracted the gene sequence (1989 bp) from the initial *Bt* genome (AMD595) using anvi’o interactive interface and used blastn v2.5.0 to blast the long-reads from MZ65. Blast hits with an alignment length > 180% of the gene length were flagged as “two copies” and hits with alignment length between 80% and 105% were flagged as “one copy”. To visualize long-reads with two copies of *BT*1871, we used minimap2 v2.17 (53) to map the MZ65 long-reads to the MZ65 genome, which had two copies of the BT1871 region. We used samtools v1.11 to extract the reads with two copies as identified above and used IGV v2.11.1 (54) to visualize the mapping and generate a figure.

### Metabolite Extraction from Cecal Material

Metabolites were extracted with the addition of extraction solvent (80% methanol spiked with internal standards and stored at -80°C, Table S4) to pre-weighed fecal/cecal samples at a ratio of 100 mg of material per mL of extraction solvent in beadruptor tubes (Fisherbrand; 15-340-154). Samples were homogenized at 4°C on a Bead Mill 24 Homogenizer (Fisher; 15-340-163), set at 1.6 m/s with 6 thirty-second cycles, 5 seconds off per cycle. Samples were then centrifuged at -10°C, 20,000 x g for 15 min and the supernatant was used for subsequent metabolomic analysis.

### Metabolite Analysis using GC-EI-MS and Methoxyamine and TMS Derivatization

Metabolites were analyzed using GCMS with Electron Impact Ionization. 100uL of metabolite extract was added to prelabeled mass spec autosampler vials (Microliter; 09-1200) and dried down completely under nitrogen stream at 30 L/min (top) 1 L/min (bottom) at 30°C (Biotage SPE Dry 96 Dual; 3579M). To dried samples, 50 **µ**L of freshly prepared 20 mg/mL methoxyamine (Sigma; 226904) in pyridine (Sigma; 270970) was added and incubated in a thermomixer C (Eppendorf) for 90 min at 30°C and 1400 rpm. After samples are cooled to room temperature, 80 **µ**L of derivatizing reagent (BSTFA + 1% TMCS; Sigma; B-023) and 70 **µ**L of Ethyl Acetate (Sigma; 439169) were added and samples were incubated in a thermomixer at 70°C for 1 hour and 1400rpm. Samples were cooled to RT and 400 **µ**L of Ethyl Acetate was added to dilute samples. Turbid samples were transferred to microcentrifuge tubes and centrifuged at 4°C, 20,000 x g for 15 min. Supernatants were then added to mass spec vials for GCMS analysis. Samples were analyzed using a GC-MS (Agilent 7890A GC system, Agilent 5975C MS detector) operating in electron impact ionization mode, using a HP-5MSUI column (30 m x 0.25 mm, 0.25 **µ**m; Agilent Technologies 19091S-433UI) and 1 **µ**L injection. Oven ramp parameters: 1 min hold at 60°C, 16°C per min up to 300°C with a 7 min hold at 300°C. Inlet temperature was 280°C and transfer line was 300°C. Data analysis was performed using MassHunter Quantitative Analysis software (version B.10, Agilent Technologies) and confirmed by comparison to authentic standards. Normalized peak areas were calculated by dividing raw peak areas of targeted analytes by averaged raw peak areas of internal standards.

## Data Availability

The data, including DNA and RNA sequencing datasets, that support the findings of this study are available in this article, the Supplemental Information, and BioProject PRJNA797447.

## Acknowledgements

We thank Hualan Liu for invaluable experimental help with the RB-TnSeq libraries. We thank David Hershey and A. Murat Eren for helpful scientific input. We also thank the Chang lab members for scientific support received.

This work was performed with support from NIH T32DK007074 (M.Z., M.K.), NIH RC2DK122394 (E.B.C.), NIH T32GM007281 (M.K.), and the Host-Microbe and Tissue and Cell Engineering cores of the UChicago DDRCC, Center for Interdisciplinary Study of Inflammatory Intestinal Disorders (C-IID) - (NIDDK P30 DK042086).

## SUPPLEMENTAL MATERIALS

**Supplemental Fig. 1:**
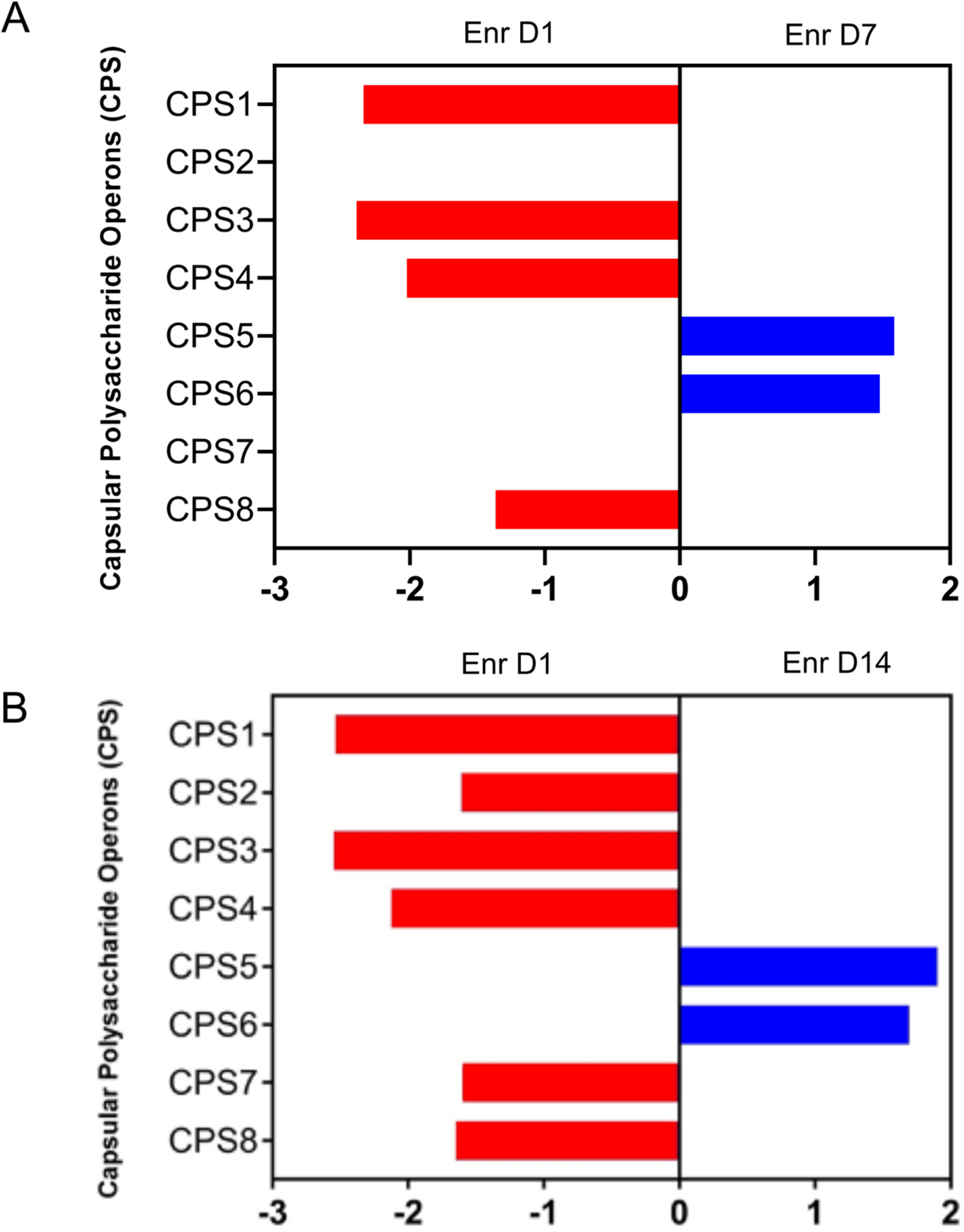
GSEA of capsular polysaccharide biosynthesis operons, comparing A) D1 to D7 expression or B) D1 to D14. Only statistically significant scores are shown (p < 0.05).

**Supplemental Fig. 2:**
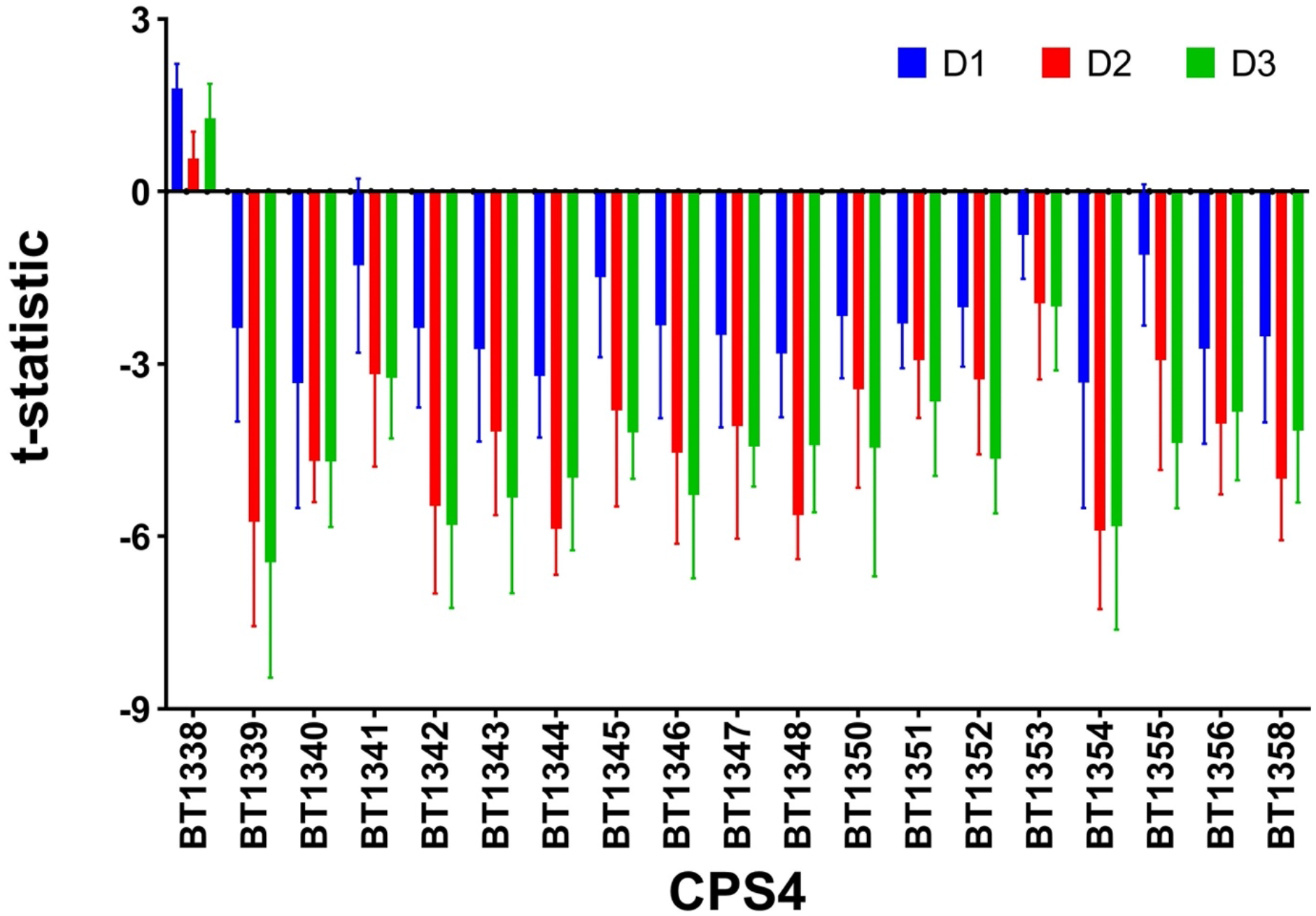
Measurement of the moderated t-statistic of Wetmore *et al*. (39) in the RB-Tn assay on all days where fitness score was measurable (D1-3) shows that gene insertions in CPS4 consistently results in statistically significant declines in mutant fitness for all genes within the CPS4 locus except for BT1338.

**Supplemental Fig. 3:**
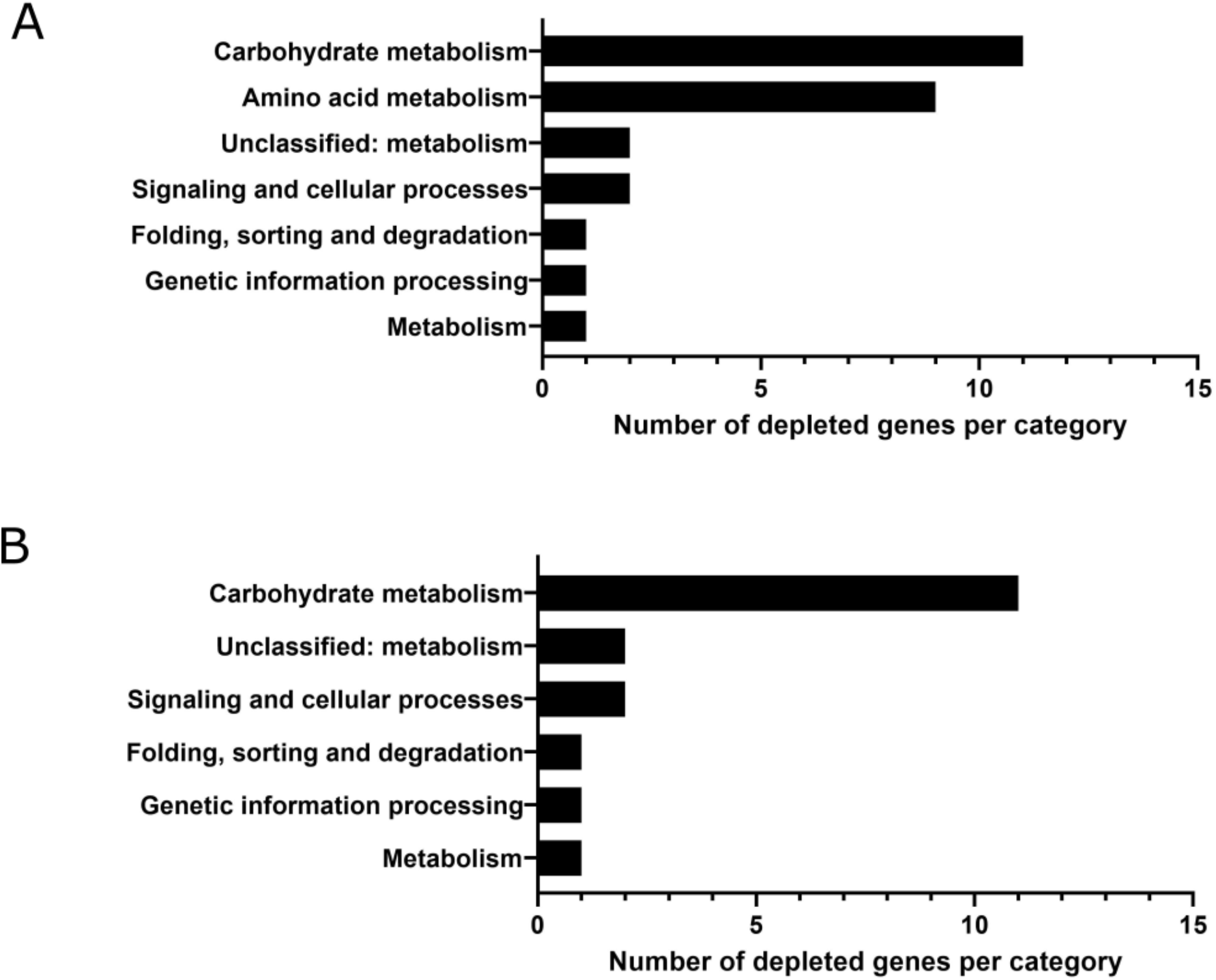
Number of genes mutants significantly depleted in the RB-Tn experiment (t-statistic < -3σ) on A) D2 or B) D3 of the experiment. Each gene was assigned a KEGG functional category if one existed in the database. See Fig. 2D for genes depleted on D1.

**Supplemental Fig. 4:**
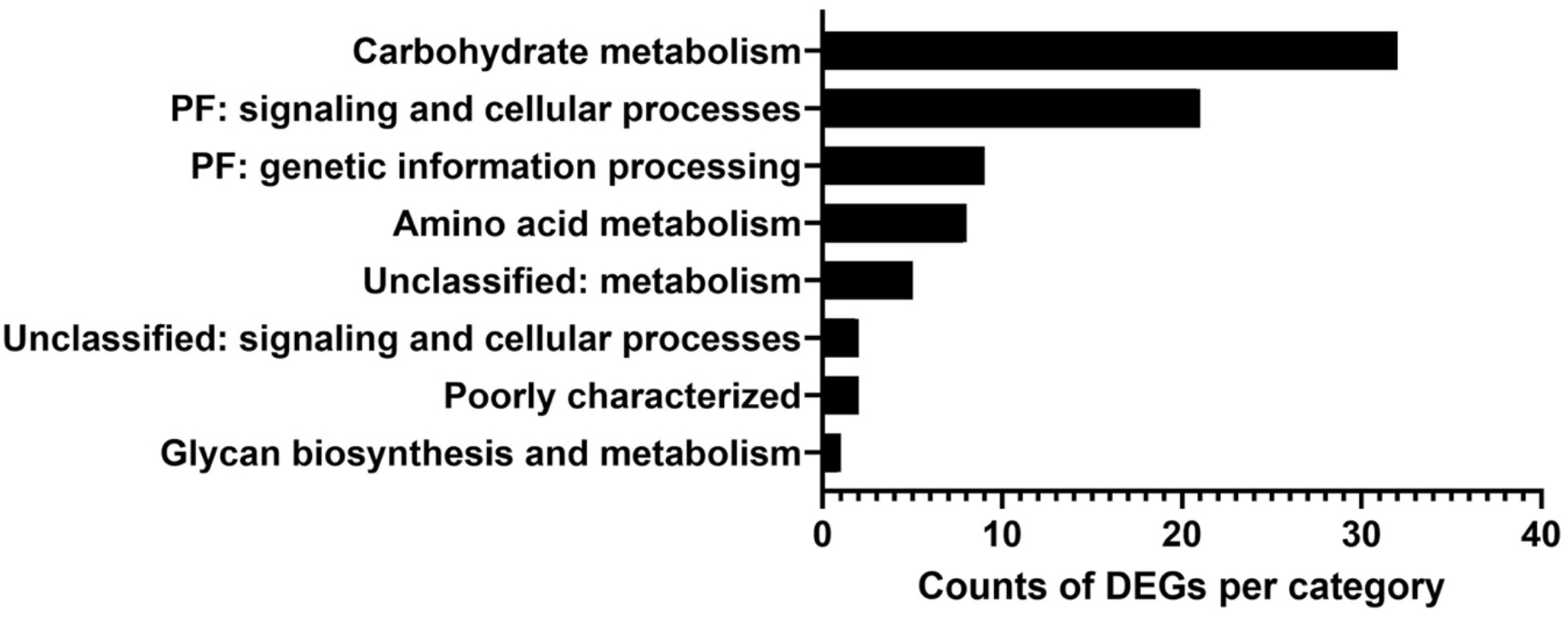
Number of DEGs enriched on D7 compared to D1 in the transcriptomics experiment [log(FDR-adjusted p-value) < -3, |log2(fold change)| > 2, and max group mean > 50 TPM]. Each gene was assigned a KEGG functional category if one existed in the database.

**Supplemental Fig. 5:**
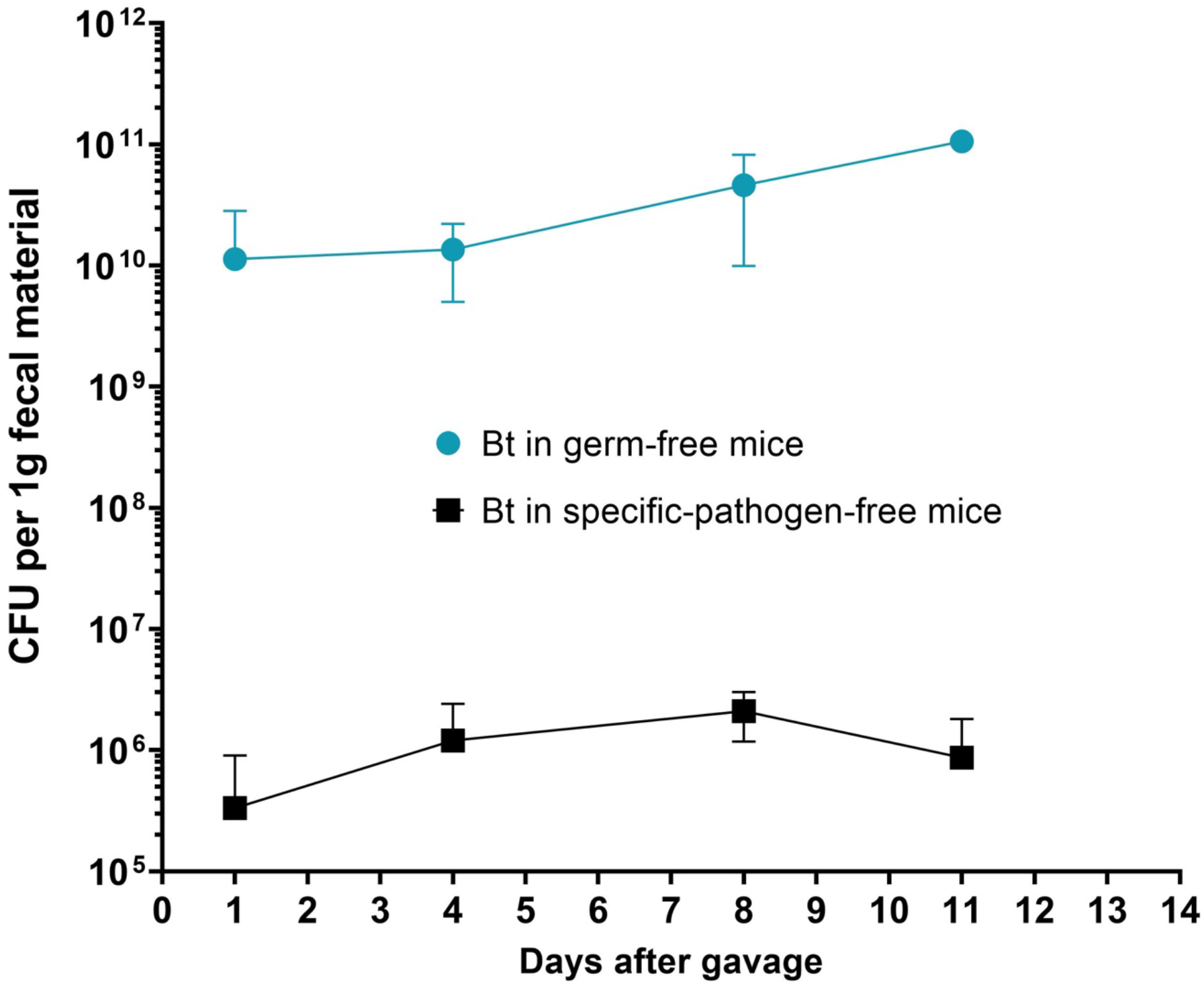
Colony forming units (CFU) for RB-Tn mutants in either germ-free (blue) or specific-pathogen-free mice (black). CFU were measured by 1/10 dilution titering on BHI-S plates and counting visible colonies after 48 hours at 37C in an anaerobic chamber. CFU is normalized for 1g of fecal material.

**Supplemental Fig. 6:**
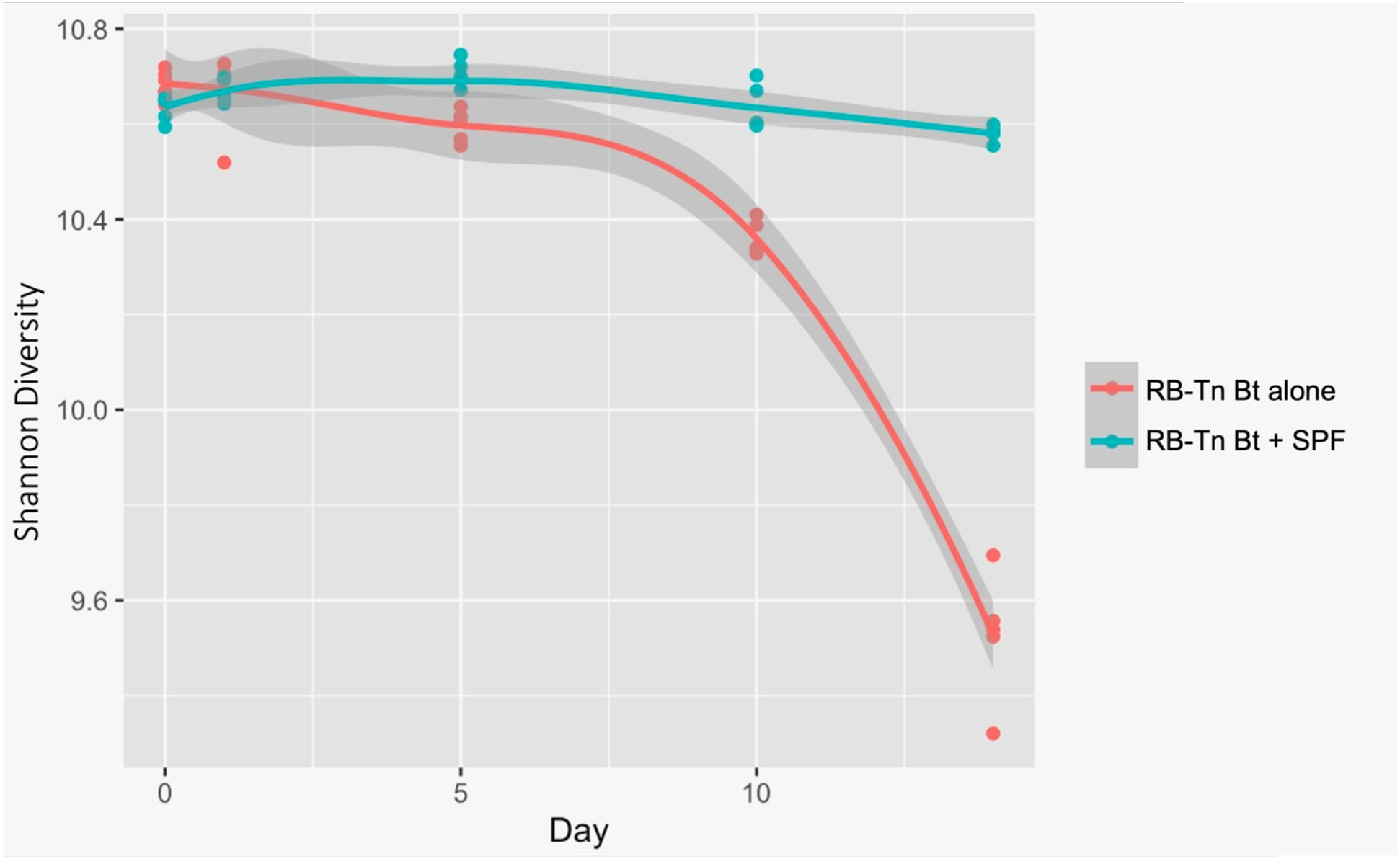
In BHI-S, Shannon Diversity of a diverse RB-Tn Bt pool decreases significantly after D4 when Bt is alone, but the Bt mutant maintain their diversity over the course of two weeks in the presence of an SPF community. Five cultures of each are maintained for 14 days with ½ dilution ever
y 12 hours.

**Supplement Fig. 7:**
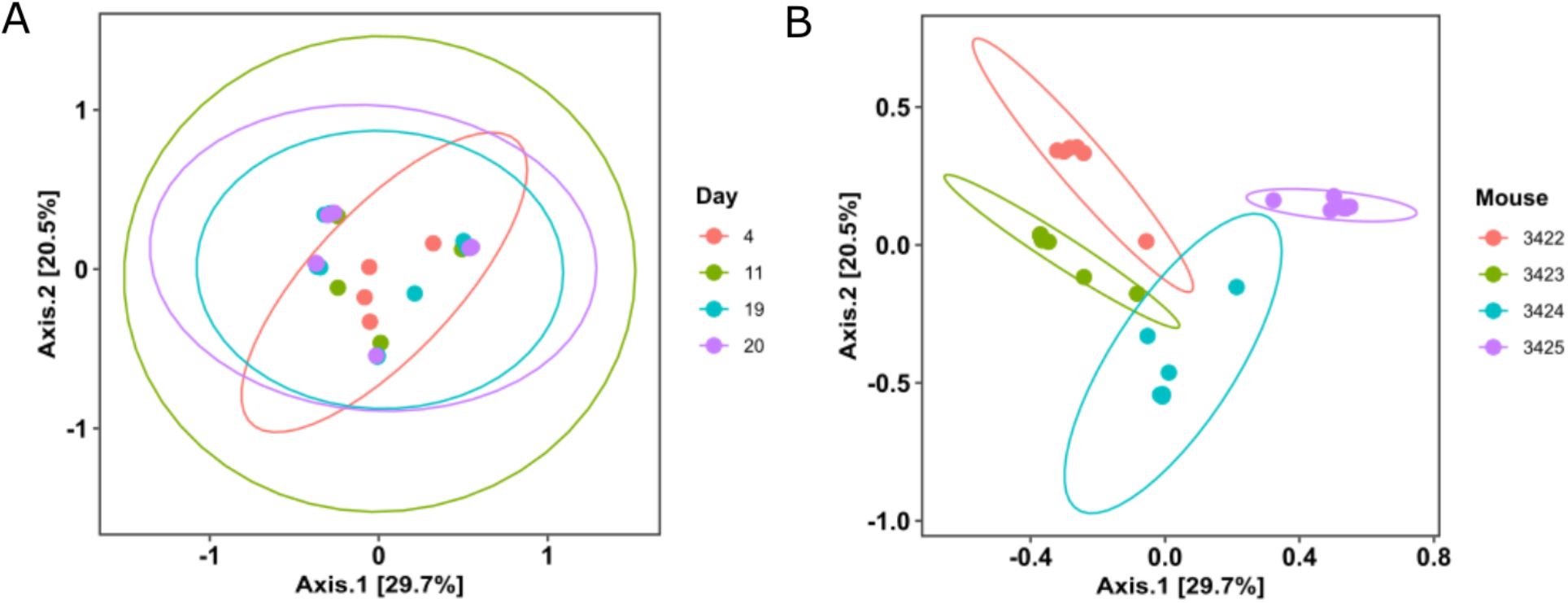
Principal Coordinates Analysis (PCoA) using Bray-Curtis dissimilarity on the relative abundance of RB-Tn mutant strains within four SPF mice, colored by A) experimental day or B) by mouse. Clustering by mouse was significant (p = 0.0001), whereas clustering by day was not.

**Supplemental Fig. 8:**
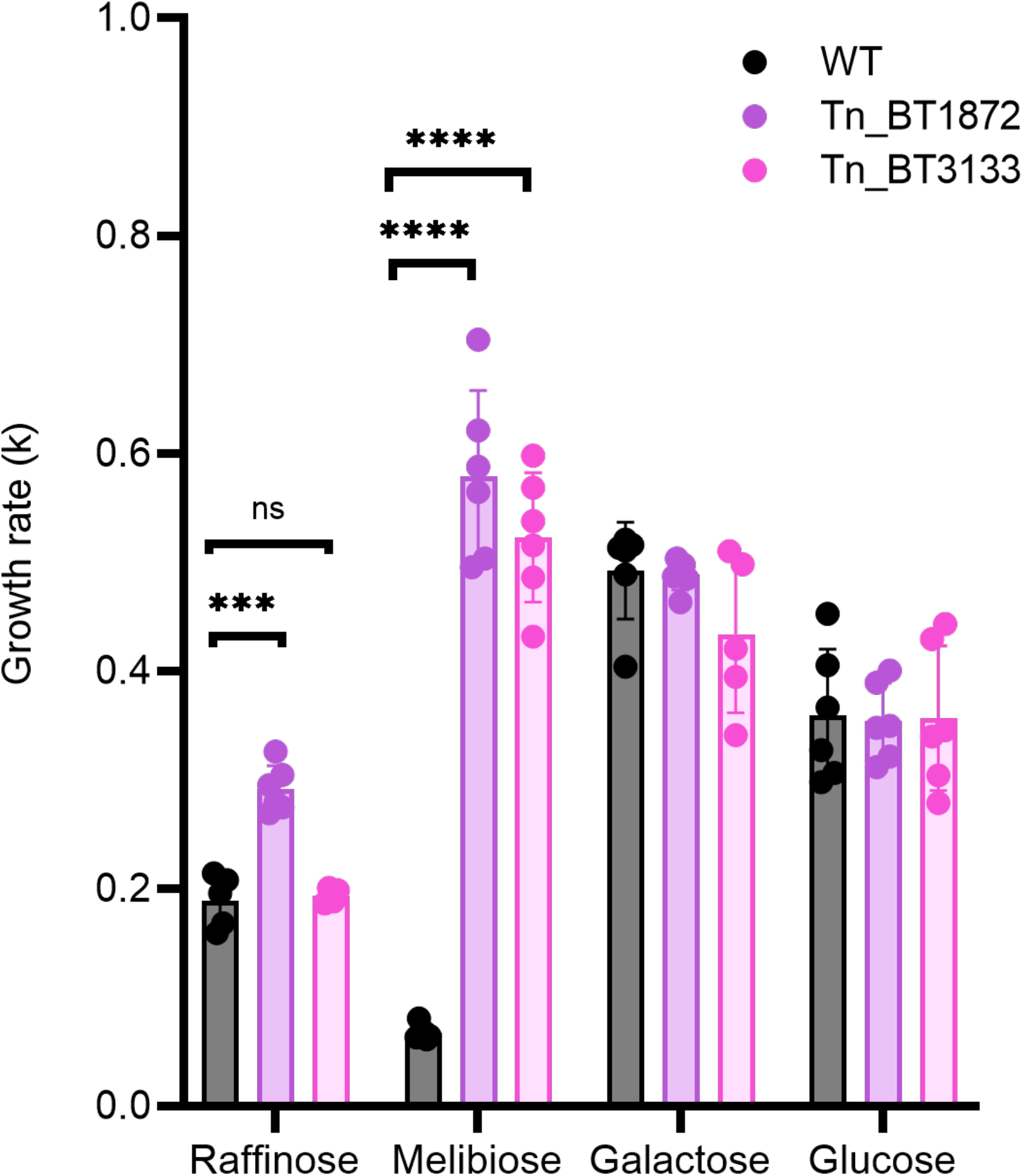
The log phase doubling times of Tn_BT3133 and Tn_BT1872 were measured in Varyl-Byrant medium with 20mM raffinose, 20mM melibiose, 20mM galactose, or 20mM glucose as the sole carbon substrate.

**Supplemental Fig. 9:**
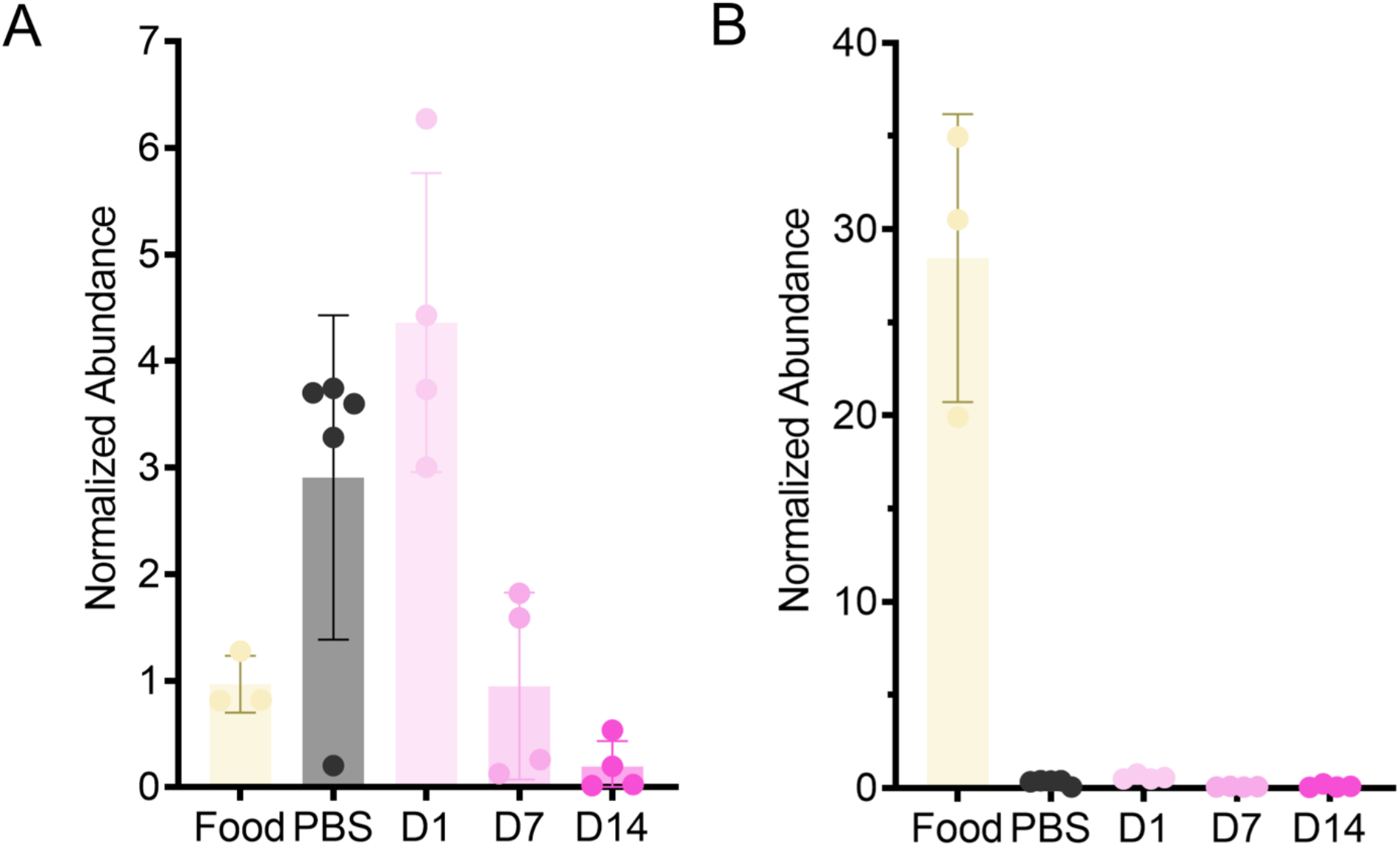
Abundance of A) melibiose or B) sucrose in the standard chow fed to GF mice, within GF ceca before colonization, or 1, 7, or 14 days post colonization.

**Supplemental Fig. 10:**
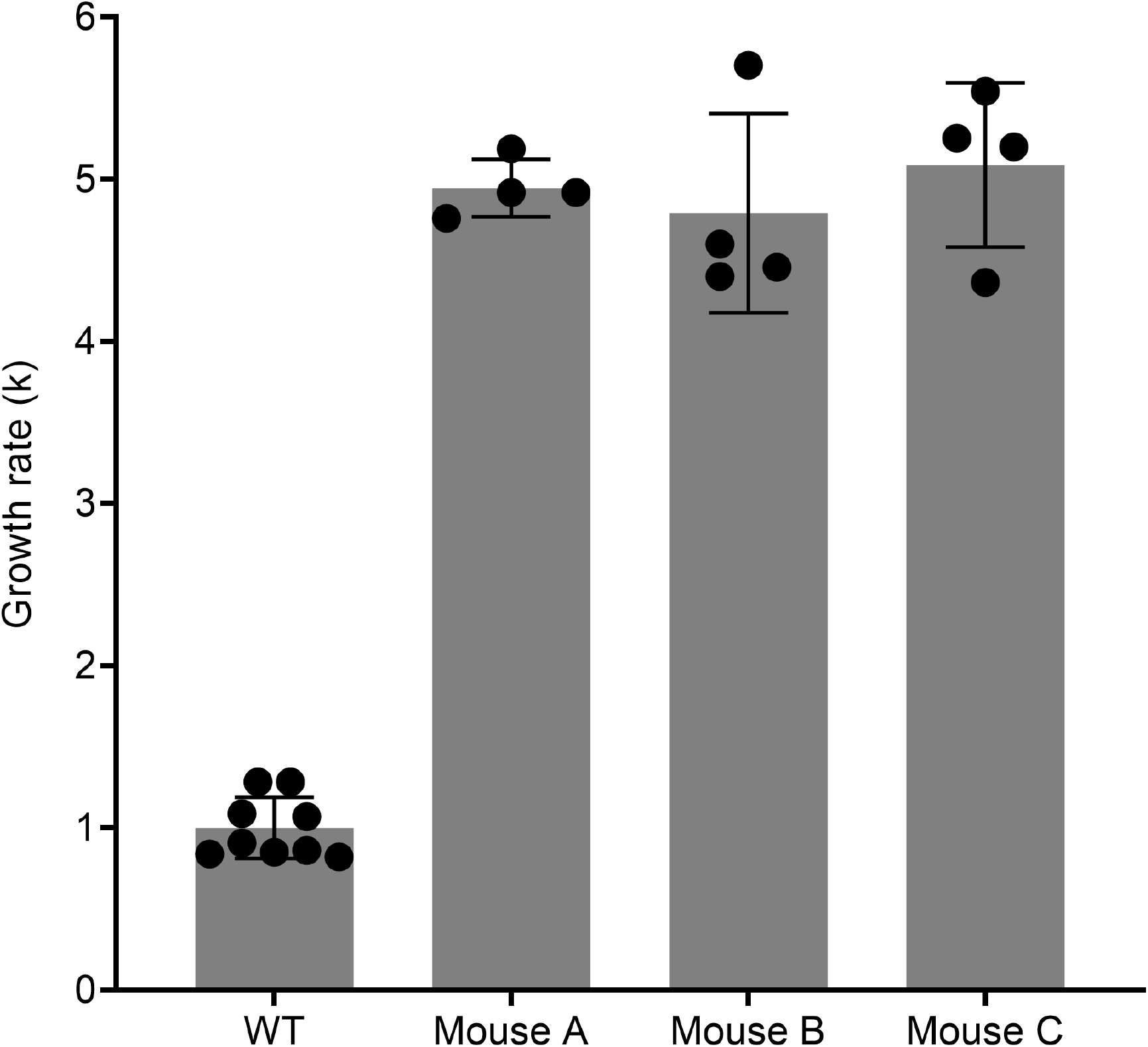
Log phase growth rate (k) of isolated strains from feces of mice colonized with WT Bt for 6 weeks compared to growth rate of WT.

**Table S1:** Strains used in this study.

| Identifier | Genotype | Source |
| --- | --- | --- |
| AMD595 | Bacteroides thetaiotaomicron VPI-5482 wildtype | Gift from Deutschbauer lab |
| AMD790 | AMD595-derived RB-Tn library | Gift from Deutschbauer lab |
| Tn_BT1874 | RB-Tn mutant isolated from monocolonized mice with barcoded transposon insertion in BT1874 | This work |
| Tn_BT1876 | RB-Tn mutant isolated from monocolonized mice with barcoded transposon insertion in BT1876 | This work |
| Tn_BT1872-83 | RB-Tn mutant isolated from monocolonized mice with barcoded transposon insertion in the intergenic region between BT1872 and BT1873 | This work |
| Tn_BT1872 | RB-Tn mutant isolated from monocolonized mice with barcoded transposon insertion in BT1872 | This work |
| Tn_BT1874/BT3132 | RB-Tn mutant isolated from monocolonized mice with barcoded transposon insertion in BT1874 and BT3132 | This work |
| Tn_BT3133 | RB-Tn mutant isolated from monocolonized mice with barcoded transposon insertion in BT3133 | This work |
| WT+BT1871OE | WT Bt with a copy of BT1871 integrated into the Bt genome at the attN1 site under the BT1311 (rpoD) constitutive promoter. | This work |
| MZ65 | Bt mutant isolated from feces after 6-week monoassociation of Bt; carries a tandem repeat of the BT1871 locus (see Fig. 7). | This work |

**Table S2:**
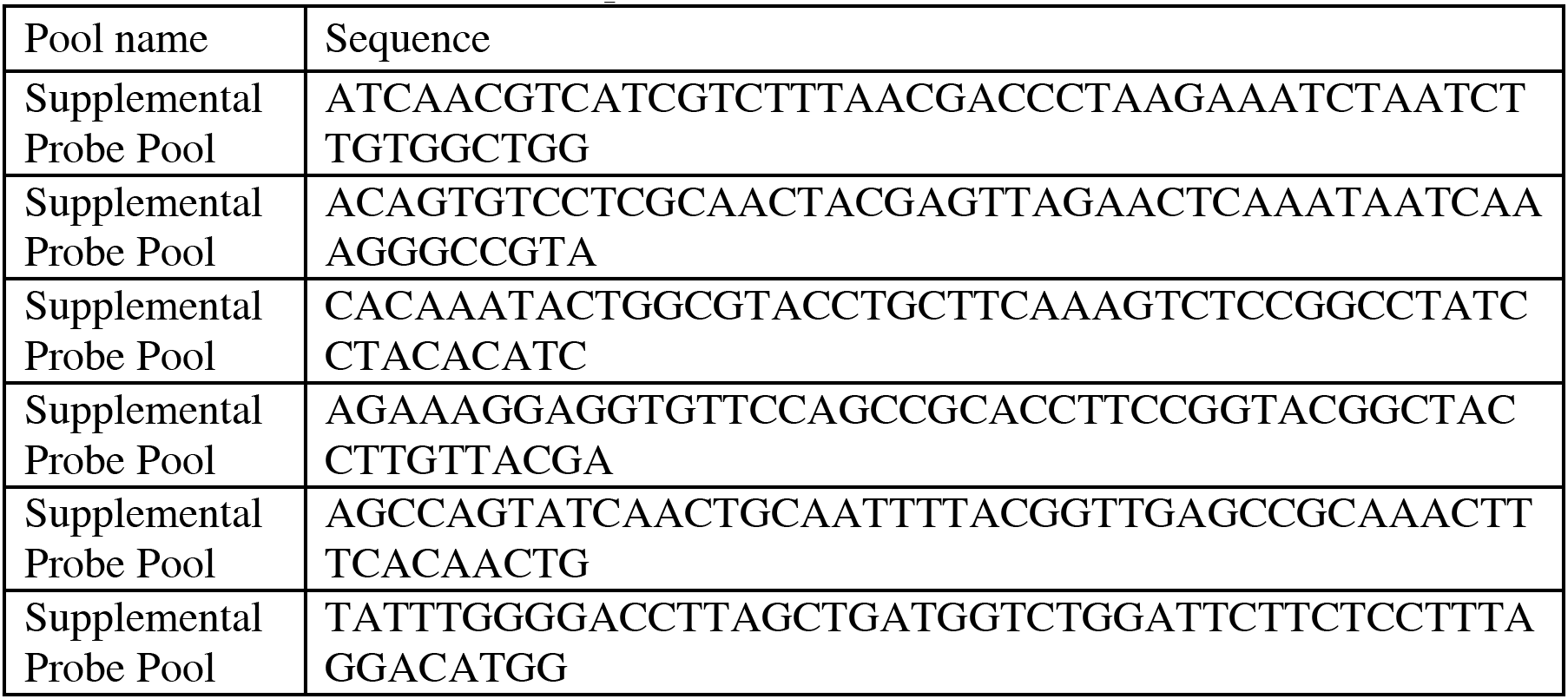
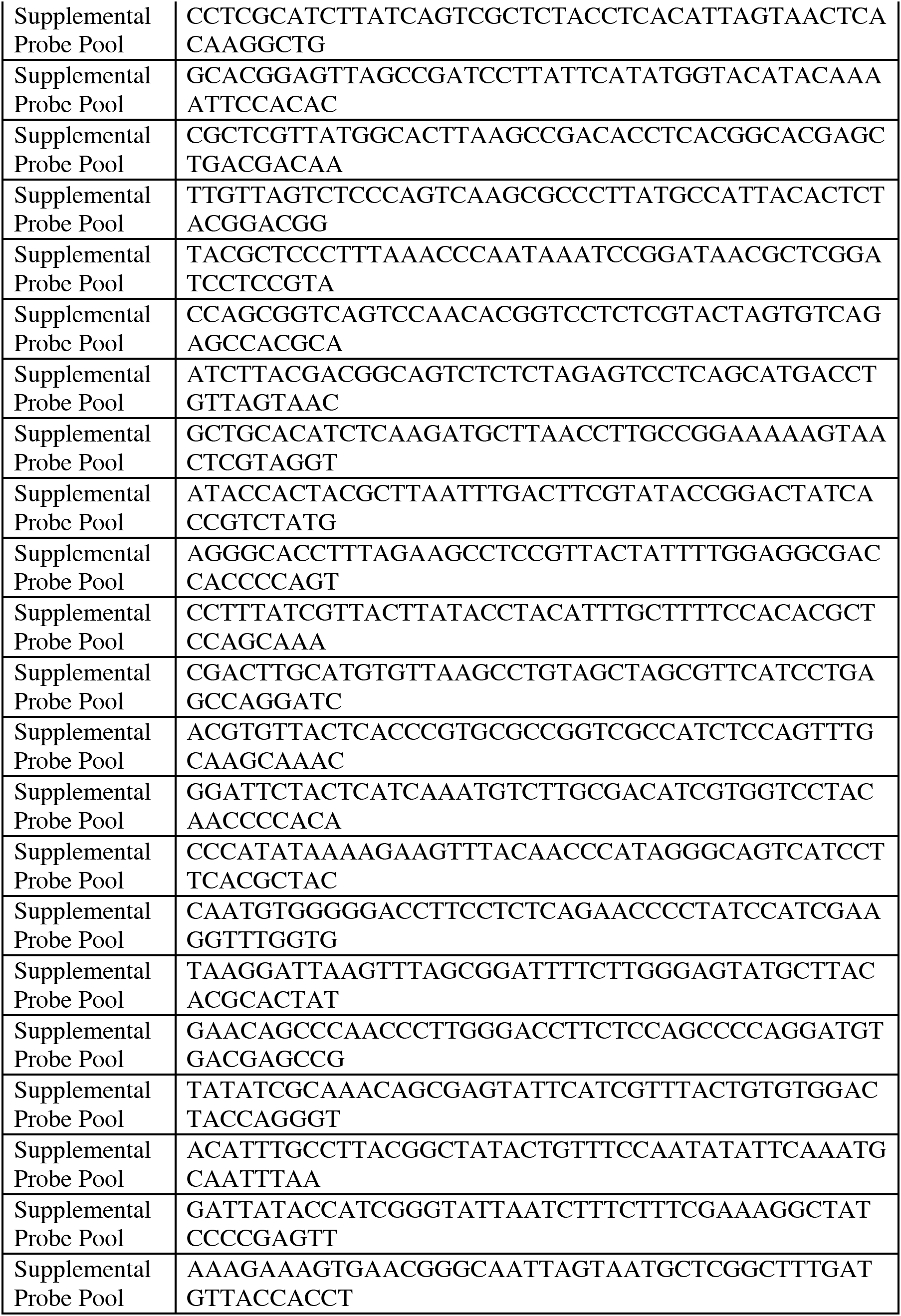
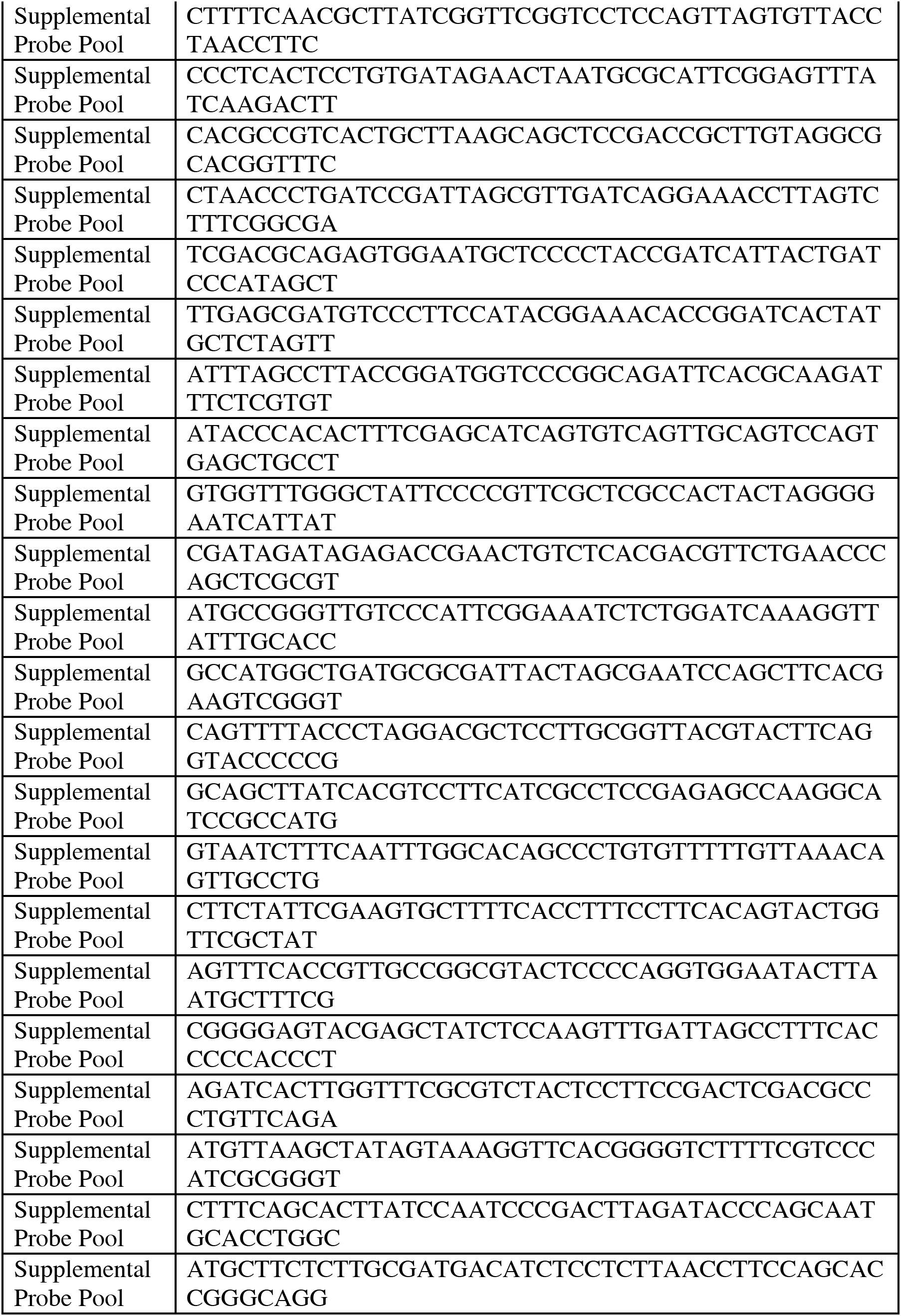
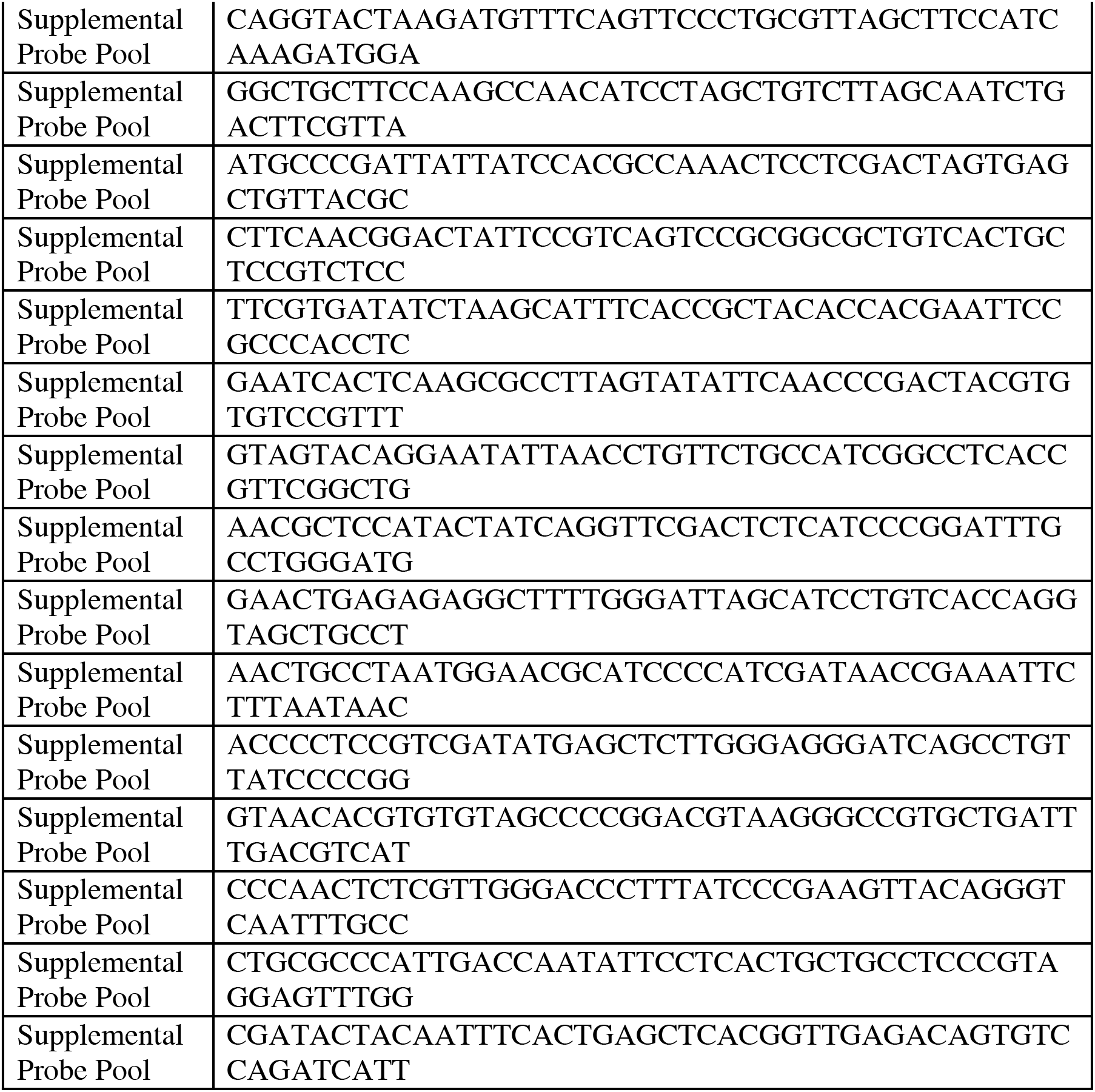
Custom Ribo-Zero Plus probes.

**Table S3:** MinION Sequencing Attributes.

|  | AMD595 | MZ55 | MZ58 | MZ65 |
| --- | --- | --- | --- | --- |
| Library Kit | SQK-RBK004 | SQK-RBK004 | SQK-RBK004 | SQK-RBK004 |
| Flowcell ID | FAQ98839 | FAQ98839 | FAQ98839 | FAQ98839 |
| Pore Type | R9.4.1 (FLO-MIN106) | R9.4.1 (FLO-MIN106) | R9.4.1 (FLO-MIN106) | R9.4.1 (FLO-MIN106) |
| Start Materials (ng) | 10,000 | 10,000 | 10,000 | 10,000 |
| DNA Shearing | 22G needle 10X, 250ul | 22G needle 10X, 250ul | 22G needle 10X, 250ul | 22G needle 10X, 250ul |
| Barcoding Input (ng) | 994.5 | 792.2 | 494.7 | 1249.5 |
| Pool Volume (ul) | 5 | 6 | 10 | 4 |
| Pool DNA Input (ng) | 497.3 | 475.3 | 494.7 | 499.8 |
| Total Pool Volume (ul) | 25 | 25 | 25 | 25 |
| Total Reads | ~1,510,000 |  |  |  |
| Total Pass Reads (%) | 1,343,227 (88.95) |  |  |  |
| Total Pass Reads per Barcode | 344,988 | 298,543 | 445,074 | 113,979 |
| Average length (bp) | 7,226 | 8,857 | 7,882 | 8,266 |
| Max. length (bp) | 108,747 | 141,745 | 114,845 | 109,375 |
| N50 (bp) | 12,596 | 16,820 | 14,970 | 15,675 |
| Top 25% Templates (bp) | 9,643 | 11,730 | 10,532 | 11,037 |
| Total Called bases | 2,493,077,325 | 2,644,475,924 | 3,508,117,512 | 942,158,294 |
| Base Calling | Guppy v5.0.11 | Guppy v5.0.11 | Guppy v5.0.11 | Guppy v5.0.11 |
| MinKNOW version | 4.3.4 | 4.3.4 | 4.3.4 | 4.3.4 |

**Table S4:** Metabolomics Internal Standards.

| Panel | ITSD | Vendor | Product Number | Concentration | Mass/Volume for 1L |
| --- | --- | --- | --- | --- | --- |
| SCFA | D3-Acetate | CIL | DLM-3126-25 | 4 mM | 340 mg |
| SCFA | D5-Propionate | CIL | DLM-1601-1 | 1 mM | 101.1 mg |
| SCFA | D7-Butyrate | CIL | DLM-7616-PK | .5 mM | 58.56 mg |
| SCFA | D9-Valerate | CIL | DLM-572-5 | .25 mM | 29.60 μL |
| SCFA | D8-Valine | CIL | DLM-488-0.25 | .05 mM | 6.26 mg |
| SCFA | D6-Succinate | CIL | DLM-831-PK | 1 mM | 124.14 mg |
| SCFA | D6-Phenol | CIL | DLM-370-PK | .025 mM | 2.5 mg |
| TMS | U13C-Palmitate | CIL | CLM-409-PK | .04 mM | 10.9 mg |
| TMS/SCFA | D7N15-Proline | CIL | DLNM-7562-0.25 | .02 mM | 2.46 mg |
| Bile Acids | D4-Cholic Acid | CIL | DLM-2611-0.05 | 1 ug/mL | 1 mL of 1 mg/mL stock |
| Bile Acids | D4-Deoxycholic Acid | CIL | DLM-2824-0.01 | 1 ug/mL | 1 mL of 1 mg/mL stock |
| Bile Acids | D4-Glycocholic Acid | CIL | DLM-2742-0.01 | 1 ug/mL | 1 mL of 1 mg/mL stock |
| Bile Acids | D4-Glycodeoxycholic Acid | CIL | DLM-9554-0.01 | .5 ug/mL | .5 mL of 1 mg/mL stock |
| Bile Acids | D4-Litchocholic Acid | CIL | DLM-9560-0.05 | 5 ug/mL | 5 mL of 1 mg/mL stock |
| Table 10: Metabolomics Internal Standards (continued) |  |  |  |  |  |
| Bile Acids | D4-Taurocholic Acid | CIL | DLM-9572-0.01 | 1 ug/mL | 1 mL of 1 mg/mL stock |
| Bile Acids | D4-Taurodeoxycholic Acid | CIL | DLM-9568-0.01 | .5 ug/mL | .5 mL of 1 mg/mL stock |
| Bile Acids | D5-Alpha-Muricholic Acid | CIL | DLM-10627-PK | 3.5 ug/mL | 3.5 mL of 1 mg/mL stock |
| Indoles | 13C11-L-Tryptophan | CIL | CLM-4290-H | 500 nM | 50 μL of 10 mM stock |
| Indoles | 13C9-L-Phenylalanine | CIL | CLM-2250-H | 500 nM | 50 μL of 10 mM stock |
| Indoles | 13C9-L-Tyrosine | CIL | CLM-2263-H | 500 nM | 50 μL of 10 mM stock |
| Indoles | D4-Serotonin | CIL | DLM-11030-0 | 500 nM | 50 μL of 10 mM stock |
| Indoles | 13C6-5-Hydroxyindole-3-acetic acid | CIL | CLM-9936-0 | 500 nM | 50 μL of 10 mM stock |
| Indoles | D4-Tryptamine | CIL | DLM-6989-0 | 500 nM | 50 μL of 10 mM stock |
| Indoles | D3-Melatonin | CIL | DLM-7101-0 | 500 nM | 50 μL of 10 mM stock |
| Indoles | 13C6-Niacin | CIL | CLM-9954-0 | 2 μM | 200 μL of 10 mM stock |
| Indoles | 13C10-L-Kynurenine | CIL | CLM-9884-0 | 500 nM | 50 μL of 10 mM stock |
| Indoles | D5-Kynurenic<br>Acid | CIL | DLM-7374-0 | 500 nM | 50 μL of 10<br>mM stock |
| Indoles | Anthranilic Acid,<br>Ring-13C6 | CIL | CLM-701-0 | 500 nM | 50 μL of 10<br>mM stock |

